# VEGFA+ macrophages promote the growth and metastasis of intrahepatic cholangiocarcinoma via OSM and THBS1 signaling

**DOI:** 10.1101/2025.10.02.679723

**Authors:** Tingjie Wang, Ruitao Long, Zhongyu Cheng, Mi Zhang, Bing Wei, Cong Lyu, Chuanrui Xu

## Abstract

Tumor-associated macrophages (TAMs) are abundant in intrahepatic cholangiocarcinoma (ICC) and correlate with poor prognosis, but their non-immunosuppressive roles remain unclear. This study aimed to define TAM transcriptional heterogeneity and map their interactions with ICC cells that promote malignancy beyond immune suppression. Single-cell and spatial transcriptome analysis with mouse and human ICC tissues revealed a TAM subpopulation, designated Mφ_VEGFA, that emerged as the major macrophage population from the intermediate stage of ICC development onward. Data from Gene Expression Omnibus (GEO) and spatial transcriptome indicate that Mφ_VEGFA was associated with poor prognosis and spatially localized to the ICC cells. Cell co-culture and cell-cell communication analyses revealed that Mφ_VEGFA promoted ICC cell proliferation via secreted OSM and downstream OSMR–STAT3–CCND1 axis, and enhanced ICC cell invasion through THBS1 and downstream CD47–mTOR/DBN1 signaling. Critically, multiplex immunofluorescence with human ICC tissue microarray indicated that *OSM* and *THBS1* were co-elevated in human ICCs, and dual blockade of both targets inhibited ICC progression in mouse models. This study uncovers the direct and critical roles of Mφ_VEGFA in driving ICC progression with paracrine signaling and further highlights the potential of targeting macrophages in ICC treatment.

## Introduction

The incidence of intrahepatic cholangiocarcinoma (ICC), which arises from the intrahepatic bile duct epithelium, is experiencing a global rise. Notably, its subtle early symptoms, aggressive nature, and chemotherapy resistance contribute to a 5-year survival rate of below 10% (1). Targeted therapies and immune checkpoint inhibitors provide benefits in some solid tumors; nevertheless, responses in ICC remain inconsistent owing to its complex tumor microenvironment (TME) (2). Tumor-associated macrophages (TAMs) are the most abundant immune cells in the ICC TME, and promote immune suppression, angiogenesis, stromal remodeling, and subsequent progression and metastasis (2, 3). Current TAM-targeting strategies, including depletion, reprogramming, recruitment control, and functional regulation, exhibit variable efficacy (4, 5). Thus, dissecting communication cascades between ICC cells and TAMs may help identify novel therapeutic targets for ICC therapy.

TAMs are highly heterogenous and distinct TAM subpopulations are reportedly associated with ICC progression. MARCO⁺ TAMs correlate with epithelial-mesenchymal transition (EMT), angiogenesis, and hypoxia in the ICC progression (6). *S100P* and *SPP1*-markers, distinguishing perihilar and peripheral cholangiocarcinoma, exhibit differential macrophage infiltration tied to prognosis and grade (7). Furthermore, spatial distribution influences TAM functions (8), and spatial-omics studies further revealed TAM heterogeneity (9, 10). Specifically, TAMs at invasive fronts mainly express pro-angiogenic factors such as VEGFA, whereas core TAMs activate immunosuppressive signaling (6). Moreover, TAM plasticity is shaped by cues such as extracellular matrix (ECM) gradients, lactate accumulation, and cytokines (9). Nonetheless, the spatial dynamics and underlying mechanisms of TAM to promote ICC development directly remain poorly defined.

Here, we integrated temporal-spatial and single-cell transcriptomics to map transcriptional landscapes, and identified a human–mouse conserved TAM population driving ICC progression. Using cell-cell communication analysis, cell-cell co-culture, multiplex immunofluorescence, and knockout mouse models, we validated the key TAM–ICC signaling axes that promote ICC proliferation and migration. These findings may clarify direct and non-immunosuppressive pro-tumoral mechanisms in TAMs, and support the development of macrophage–targeted immunotherapy for ICC.

## RESULTS

### Mφ_VEGFA is enriched in ICC and required for ICC progression

To characterize macrophage subsets involved in ICC malignancy, we established an AKT1/NICD1-induced ICC mouse model via hydrodynamic tail vein injection. Histological examination of mouse ICC tissues collected at four time points revealed progressive tumor development (Supplementary Figure S1A). Tumor budding was first observed on day 7, followed by the formation of ductal-like structures by day 15 and the development of more defined nodules with disorganized structures by day 25. By day 31, ICC tumor nodules with bile-like fluid had spread extensively across the liver surface, indicating advanced-stage disease (11). Therefore, we defined ICC tissues collected on day 7 as early stage, on day 15 as middle, on day 25 as middle-advanced, and on day 31 as advanced.

scRNA-seq was performed using tissues collected at all stages, and nine major cell types, including macrophages, were identified (Supplementary Figure S1B–C). Subsequent re-clustering analysis identified seven macrophage subtypes: Mφ_Vegfa, Mφ_Dnajb1, Mφ_S100a4, Mφ_Chil3, Mφ_Mki67, Mφ_Clec4f, and Mφ_Cxcl9 (Figure 1A). In the early and middle stages of ICC progression, Mφ_Cxcl9 and Mφ_S100a4 showed dominant infiltration (Figure 1B). Mφ_Cxcl9 expressed high levels of *Cxcl9* and *Cxcl10*, whereas Mφ_S100a4 expressed high levels of *S100a4* and *S100a6* (Supplementary Figure S1D). In addition, these two macrophage subtypes demonstrated enriched pathways associated with antitumor immune response (Supplementary Figure S1E). Of note, Mφ_Clec4f exhibited high levels of tissue-resident markers *Vsig4* and *Clec4f*, infiltrated highly at all time points except day 25, and mediated tissue repair and homeostasis, suggesting that it represents a Kupffer-like cell type (Figure 1B and Supplementary Figure S1D)(12). In the middle**–**to**–**advanced or advanced stages, Mφ_Chil3, Mφ_Dnajb1, and Mφ_Vegfa were markedly infiltrated (Figure 1B). Specifically, Mφ_Chil3 highly expressed *Chil3*, *S100a4*, *S100a6,* and *S100a8,* and were enriched pathways associated with inflammation (Figure 1B and Supplementary Figure S1D). Mφ_Dnajb1 highly expressed *Hspb1*, *C1qa* and *Dnajb1*, whereas Mφ_Vegfa expressed canonical M2-like markers, including *Vegfa*, *Gpnmb*, *Trem2*, *Spp1* and *Arg1* (Supplementary Figure S1D) (13, 14). Notably, both Mφ_Dnajb1 and Mφ_Vegfa showed enriched pathways related to stress responses and cellular signaling/communication (Supplementary Figure S1E). Critically, those macrophage subtypes except Mφ_Chil3 and Mφ_Dnajb1 identified in AKT1/NICD1 mouse ICC tissues were also identified in AKT1/YAP1 mouse ICC tissues using publicly available scRNA-seq data (15) (Supplementary Figure S2A-B).

**Figure 1:**
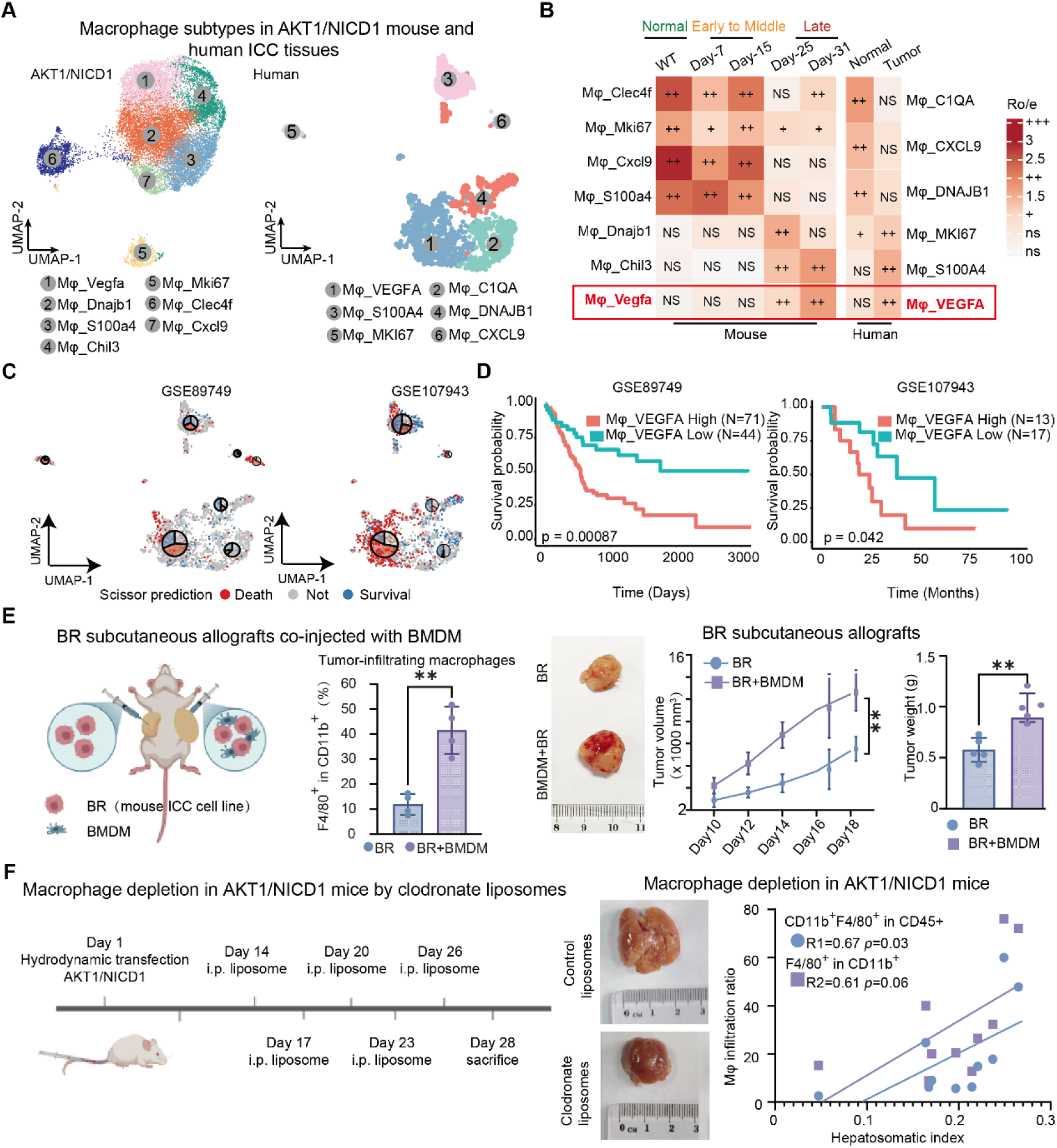
Mφ_VEGFA is enriched in ICC and promotes ICC progression in mice. **(A)** Macrophage distribution in AKT1/NICD1 mouse and human tissues. The UMAP plot visualizes macrophage cell types. **(B)** Temporal distribution of macrophage subtypes in AKT1/NICD1-induced mouse ICC. **(C)** Scatter-pie and plots illustrate prognostic associations of distinct macrophage subtypes calculated by Scissor. The pie chart illustrates the association between patient prognosis outcomes and each macrophage subtype. Blue, survival; red, death; grey, no significant association. **(D)** Prognosis of Mφ_VEGFA-high and Mφ_VEGFA-low patients with ICC in two bulk RNA data sets (GSE89749 and GSE107943). Statistical significance was calculated using the log-rank test. **(E)** Allografts established by co-injection of bone marrow-derived macrophages (BMDMs) and mouse ICC cells (BR cells) in a 1:1 ratio in mice. From left to right: schematic illustration of allograft model, quantification of tumor-infiltrating macrophages (F4/80⁺CD11b⁺ cells) (n = 4 mice per group), representative images of allografts, tumor volume growth curve (n = 6 mice per group), and tumor weights (n = 6 mice per group). Data are presented as mean ± SD; Significance was determined by a two-tailed unpaired Student t-test. *\*\* P* < 0.01, and n = 4 mice. **(F)** Macrophage depletion in the AKT1/NICD1 ICC mouse model. The control group mice received blank liposomes. Macrophage infiltration levels (R1, CD11b⁺F4/80⁺ cells within CD45⁺ population; R2, F4/80⁺ cells within CD11b⁺ population) were analyzed via flow cytometry. Correlation with liver-to-body weight ratio was analyzed using Spearman’s rank correlation test. Abbreviations: ICC, intrahepatic cholangiocarcinoma; NICD1, NOTCH1 intracellular domain; BR cells, cells from BMI1/NRAS-driven primary ICC tissues in FVB mice; R1, R2, correlation coefficient.

To further validate these findings in human ICC, we integrated two human ICC scRNA-seq datasets, GSE138709 (16) and GSE189903 (17) (Supplementary Table S2), and identified six similar macrophage subtypes (Figure 1A, Supplementary Figure S2C). Notably, the human macrophage subpopulations Mφ_CXCL9 and Mφ_VEGFA displayed highly similar gene expression profiles, enriched pathways, and tissue-infiltration patterns to the corresponding mouse macrophage subpopulations Mφ_Cxcl9 and Mφ_Vegfa (Figure 1B, Supplementary Figure S2C–D). Furthermore, Mφ_DNAJB1 and Mφ_S100A4 exhibited similar gene expression profiles and enriched pathways to Mφ_Dnajb1 and Mφ_S100a4, respectively (Supplementary Figure S2C–D). However, unlike the patterns observed in mouse ICC tissues, Mφ_DNAJB1 was predominantly enriched in normal tissues, whereas Mφ_S100A4 was enriched in malignant tissues in human ICC samples. In contrast, the distribution of Mφ_CXCL9 and Mφ_VEGFA demonstrated strong concordance between mouse and human ICC tissues. Mφ_CXCL9 mainly infiltrated into adjacent tissues, and Mφ_VEGFA was dominant in advanced-stage mouse and human malignant tissues (Figure 1B). Together, we hypothesized that Mφ_VEGFA is associated with ICC progression among these macrophage subtypes.

To test this hypothesis, we then analyzed the correlation between different macrophage subtypes and ICC prognosis. Using the Scissor algorithm (18), we found that Mφ_VEGFA among the six macrophage subtypes showed the strongest association with mortality in patients with ICC in the GSE89749 (19) and GSE107943 (20) datasets (Figure 1C, Supplementary Table S2). Next, we selected the top 10 signature genes specifically enriched in Mφ_VEGFA and performed survival analyses (Supplementary Table S4). Patients with higher levels of Mφ_VEGFA-associated gene signatures had significantly worse outcomes than those with lower levels of signature gene expression, further confirming the potential roles of Mφ_VEGFA in the malignant progression of ICC (Figure 1D).

We then conducted *in vivo* experiments to validate the function of macrophages in ICC development. Mouse ICC allografts were established by injecting BR ICC cells (isolated from BMI1/NRAS mouse ICC tissues) with or without bone marrow-derived macrophages (BMDMs). As expected, co-injection of BR cells and BMDMs significantly accelerated tumor growth compared to injection of BR cells alone (Figure 1E, Supplementary Figure S3A). Flow cytometry analysis of allografts isolated on Day 18 after tumor inoculation revealed increased infiltration of F4/80⁺CD11b⁺ macrophages in BR/BMDM co-injected allografts relative to BR allografts (Figure 1E, Supplementary Figure S3B–D). Then, macrophage depletion was performed by injecting clodronate liposome intraperitoneally every three days starting on day 14 after hydrodynamic tail-vein injection of AKT1/NICD1 in mice (Figure 1F). Macrophage depletion suppressed tumor progression, as evidenced by reduced ICC tumor burden in clodronate liposome-treated mice. Furthermore, macrophage infiltration levels (CD11b^+^F4/80^+^ within CD45^+^ and F4/80^+^ within CD11b^+^) positively correlated with the liver-to-body weight ratio (Figure 1F, Supplementary Figure S4). Together, these findings indicate that Mφ_VEGFA displays M2-like features and its infiltration is strongly associated with the malignant progression of ICC.

### Mφ_VEGFA is localized adjacent to ICC cells and promotes ICC progression through cytokine secretion

Considering the strong association between Mφ_VEGFA and ICC malignancy, we then examined the detailed localization of Mφ_VEGFA in human ICC tissues using multiplex immunofluorescence (mIF). Notably, a pronounced fluorescence signal of VEGFA+CD68^+^ macrophages indicated their localization or penetration with a proximity to the ICC cells majorly around 0**–**20 µm in two independent human ICC tissue arrays, indicating close localization between Mφ_VEGFA and ICC cells (Figure 2A). Next, using high-resolution 10x Visium HD assay and the BANKSY method (21), we found seven major cell types in human ICC tissues (Figure 2B). Mφ_VEGFA showed strong co-occurrence with ICC cells and could serve as a predictor for CK19^+^ ICC cells based on mistyR (22) (Figure 2B, Supplementary Figure S5A). Using 10x Visuim data from human ICC tissue (23), we segmented the human ICC tissues into non-malignant, boundary, and malignant regions (Figure 2C). Mφ_VEGFA cell type scores exhibited a spatial increase from non-malignant to malignant regions. Pathway enrichment analysis showed that the cell cycle pathway from the KEGG and epithelial-to-mesenchymal transition (EMT) pathway from the GO database (Biological Process, BP) were most enriched in the malignant region (Figure 2D). Importantly, spatial co-occurrence between Mφ_Vegfa and ICC cells was confirmed in AKT1/NICD1 mouse ICC tissues using the BMKMANU S1000 ST assay (Supplementary Figure S5B). Together, these results suggest that Mφ_VEGFA is highly co-localized with ICC cells and may favor ICC progression.

**Figure 2:**
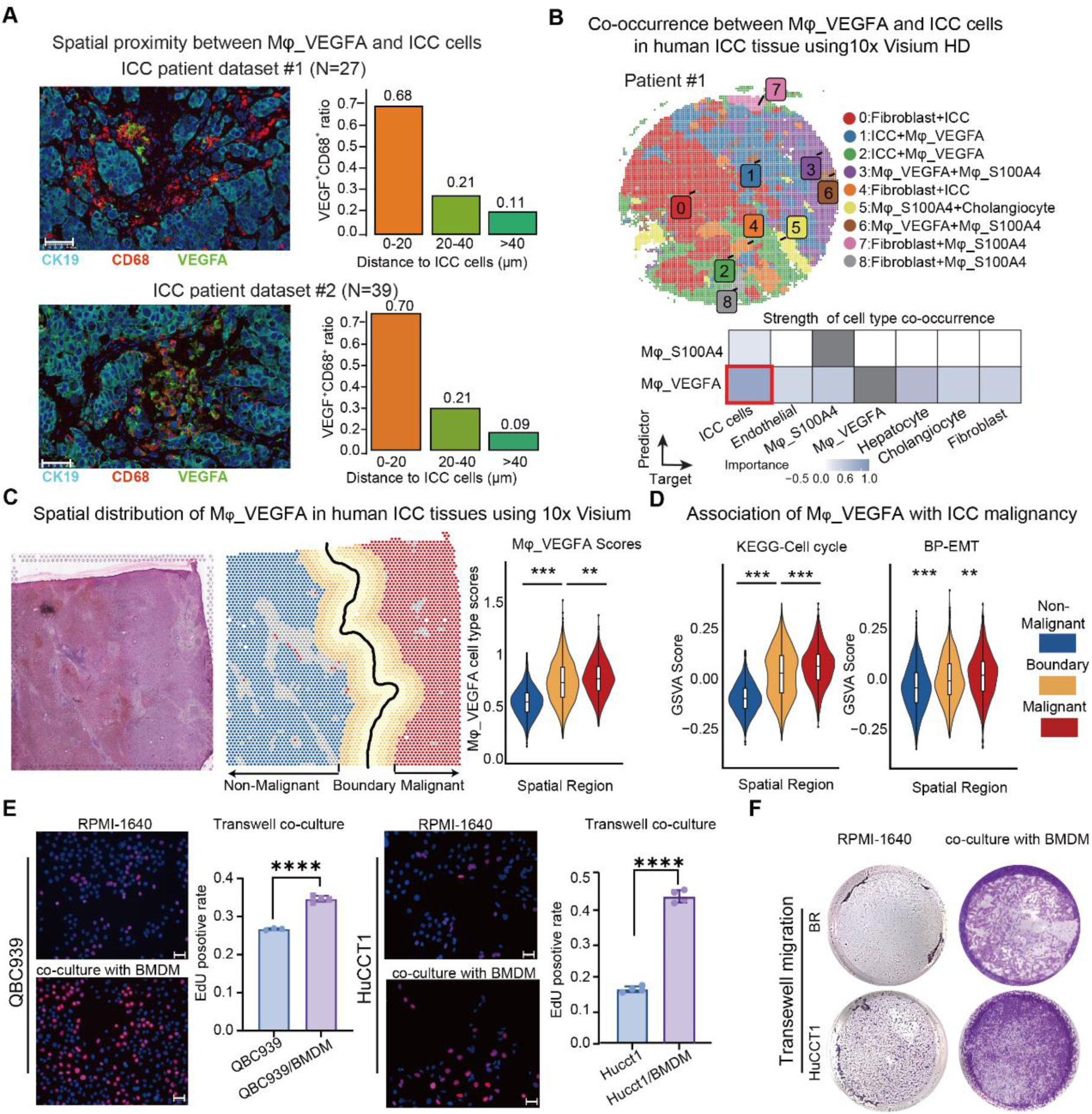
Mφ_VEGFA is spatially proximal to ICC cells and promotes ICC malignant progression. **(A)** Multiplex immunofluorescence (mIF) showing the spatial distribution of VEGFA^+^CD68^+^ macrophages and ICC cells (CK19^+^) and quantification of the proportion of VEGFA^+^CD68^+^ macrophages according to their distance to ICC cells. **(B)** Spatial domains defined by BANKSY in human ICC samples using the 10x HD platform (upper) and significance of cell type abundance in predicting the abundance of other cell types in the local niche (lower). **(C)** H&E staining of the calculated spatial distribution of Mφ_VEGFA cells in human ICC tissues. **(D)** Violin plots showing the distribution of malignant cells according to proliferation and epithelial-mesenchymal transition scores. **(E)** EdU incorporation assays in QBC939 and HuCCT1 cells cultured alone or with BMDMs in a Transwell system. Scale bar = 50 μm. **(F)** Transwell migration assays for BR and HuCCT1 cells cultured alone or co-cultured with BMDMs. Abbreviations: H&E, Hematoxylin and Eosin staining. Data are presented as mean ± SD. Significance was determined by a two-tailed unpaired Student t-test (*\*\*\*P* < 0.001, *\*\*\*\* P* < 0.0001, and n = 4).

Next, we investigated the stimulatory roles of Mφ_VEGFA on ICC cells using multiple cell-cell co-culture systems (Supplementary Figure S6A). Non-contact Transwell co-cultures of RAW264.7 cells or BMDMs with QBC939 or BR-derived tumor-conditioned medium generated TAMs with robust upregulation of *VEGFA* and M2 markers, including *SPP1*, *ARG1*, *VEGFA*, *IL10*, and *CD206* (Figure S6B–F). EdU incorporation and Transwell assays demonstrated that macrophage co-culture significantly enhanced ICC cell proliferation and migration, respectively (Figure 2E–F). Therefore, VEGFA+ macrophages may promote ICC cell proliferation and migration in a contact-independent manner.

### Mφ_VEGFA promotes ICC proliferation via the OSM-OSMR-STAT3 axis

To explore mechanisms underlying Mφ_VEGFA-mediated ICC progression, we re-clustered epithelial cells from AKT1/NICD1 mouse ICC tissues and obtained eight epithelial cell subpopulations (Figure 3A, Supplementary Figure S7A–B). Among these populations, E_Alb was defined as hepatocytes and E_Fxyd2 as cholangiocytes, supported by the lowest CNV scores consistent with non-malignancy (Figure 3A). Conversely, E_Jun and E_Cd44 demonstrated the highest CNV scores and were enriched for proliferation-associated pathways, including cell cycle, cell proliferation, and MAPK cascade, as well as invasion- and metastasis-related signaling, including mesenchyme development and cell motility (Figure 3A). Therefore, E_Jun and E_Cd44 represent ICC cell populations with high proliferative capacity and malignancy. Next, we integrated three cell–cell communication approaches (CellChat, CellphoneDB, and NicheNet) to analyze interactions between Mφ_VEGFA and the ICC cell populations E_Jun and E_Cd44 in mice (Figure 3B). Downstream signaling pathway analysis indicated that Mφ_VEGFA engages ICC cells via OSM and THBS1 ligands (Figure 3B). Further analysis demonstrated that *Osm* interacts with E_Cd44/E_Jun cells via the receptors *Osmr* or *Lifr* (Figure 3C, Supplementary Figure S7C). scRNAseq data from human ICC tissues also confirmed macrophage–tumor communication via OSM/OSMR (Figure 3D, Supplementary Figure S7D). Spatial interaction analysis and spatial mapping by mIF further confirmed co-localization of OSM⁺ macrophages with OSMR⁺ ICC cells (Figure 3E, Supplementary Figure S7E–G).

**Figure 3.**
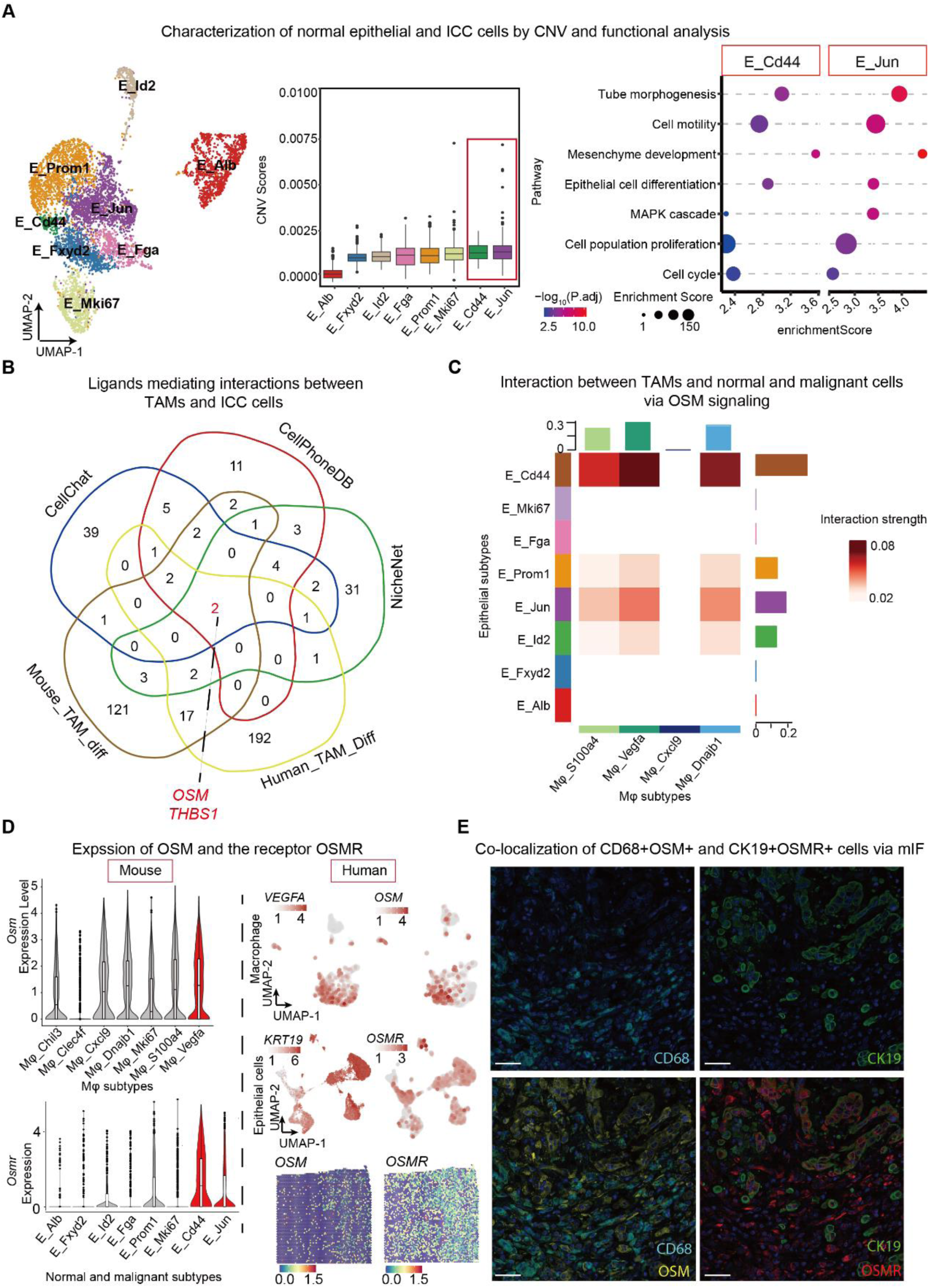
Mφ_VEGFA interacts with ICC cells via OSM-OSMR. **(A)** UMAP (left) and box plots (middle) illustrate the CNV scores of epithelial subtypes in AKT1/NICD1. Bubble chart visualizing pathway enrichment results for the subtypes E_Jun and E_Cd44 with the highest CNV scores (right). **(B)** Venn diagram demonstrating enriched ligands significantly upregulated in both Mφ_Vegfa (mouse, Mouse_TAM_diff) and Mφ_VEGFA (human, Human_TAM_Diff) identified by three cell communication methods (CellphoneDB, CellChat, and NicheNet). **(C)** Heat map showing the interaction intensity of *OSM* signaling between macrophage and ICC cells in AKT1/NICD1 mouse ICC tissues. **(D)** Violin plots show *Osm* in macrophage and its receptor *Osmr* expression in epithelial subtypes in mouse ICC tissues (left). Scatter plots show the OSM-OSMR gene pair in human ICC single-cell and spatial RNA-seq data (right). **(E)** mIF showing spatial co-localization between CD68⁺OSM⁺ TAMs and OSMR⁺KRT19⁺ ICC cells in human ICC tissues. Scale bar, 200 μm.

To identify the downstream signaling of OSM/OSMR and LIFR, we used NicheNet software to predict target genes for the ligands regulated by OSM for further analysis. These genes were enriched in proliferation-related pathways, including JAK-STAT, PI3K-AKT, mTOR, and cell cycle pathways (Figure 4A). To test whether *OSMR* and *LIFR* are required for ICC proliferation, we generated *Osmr*- or *Lifr*-liver conditional knockout mice Osmr^fl/fl^; Alb-Cre and Lifr^fl/fl^; Alb-Cre, and hydrodynamically injected them with AKT1/NICD1 or AKT1/YAP1 to induce ICC (Figure 4B). Compared with Lifr^fl/fl^ mice, Lifr^fl/fl^; Alb-Cre mice demonstrated no significant difference in liver-to-body weight ratio upon AKT1/NICD1 injection (Figure 4C). Nevertheless, Osmr^fl/fl^; Alb-Cre mice exhibited a marked reduction of liver-to-body weight ratio after AKT1/NICD1 or AKT1/YAP1 injection compared with that in Osmr^fl/fl^ mice (Figure 4C, Supplementary Figure S8A). These results indicate that OSMR, but not LIFR, mediates communication between Mφ_VEGFA and ICC cells.

**Figure 4.**
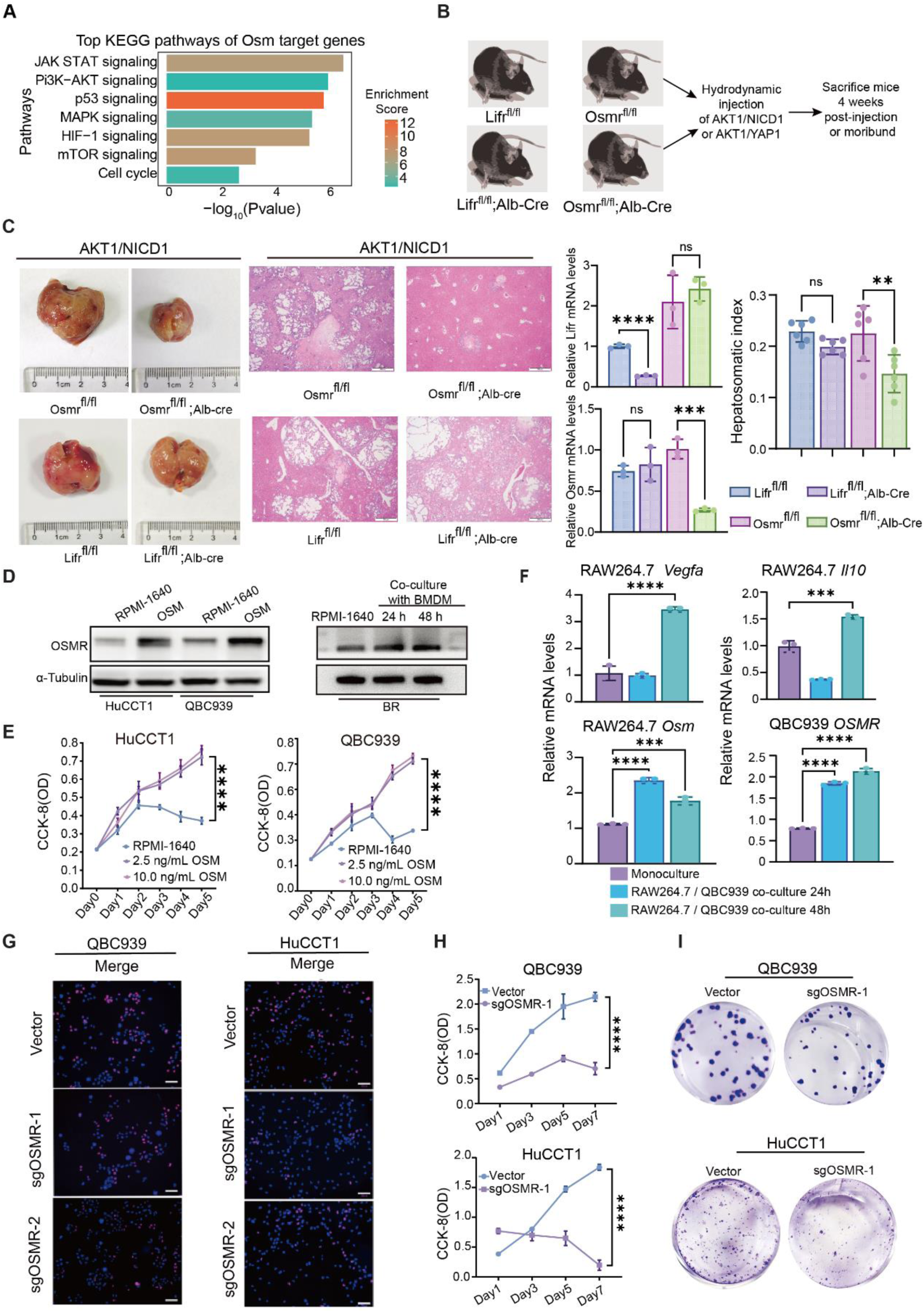
Mφ_VEGFA promotes the proliferation of ICC tissue and cells via OSM-OSMR. **(A)** Bar plot showing the top KEGG pathways of Osm target genes obtained by NicheNet. **(B and C)** Schematic illustration of experimental procedure and representative images of livers from Lifr^fl/fl^; Alb-Cre, Osmr^fl/fl^; Alb-Cre, and corresponding control mice injected with AKT1/NICD1. Scale bar, 200 μm. **(D and E)** Western blot and CCK-8 assay for cell viability measurements of HuCCT1 and QBC939 cells treated with 2.5 ng/mL or 5.0 ng/mL OSM for 5 days. **(F)** Relative mRNA levels of VEGF, IL10, and OSM in RAW264.7 or OSMR in QBC939 cells upon co-culture. **(G)** EdU incorporation assay in QBC939 cells transfected with two OSMR sgRNAs. Scale bar, 20 μm. **(H)** CCK-8 assay measuring the growth of HuCCT1 and QBC939 cells with sgOSMR-1 transfection at days 1, 3, and 5 after seeding. **(I)** Colony formation assay of HuCCT1 and QBC939 cells transfected with sgOSMR-1. Statistical significance was analyzed using one-way ANOVA (*P* < 0.05), followed by Dunnett’s t-test (ns, not significant, ** *P* < 0.01, *** *P* < 0.001; **** *P* < 0.0001).

Recombinant OSM treatment or co-culture with BMDMs increased OSMR levels in ICC cells and promoted ICC cell growth (Figure 4D**–**E, Supplementary Figure S7B). Similarly, RAW264.7 and QBC939 co-cultures increased *OSM* mRNA levels in macrophages and increased OSMR mRNA levels in QBC939 cells (Figure 4F, Supplementary Figure S7C). OSMR knockdown with siRNA suppressed ICC cell viability (Supplementary Figure S8D–E). Likewise, knockout of OSMR with sgRNA significantly decreased ICC cell proliferation, viability, and clonogenicity (Figure 4G–I, Supplementary Figure S7F).

We then analyzed downstream targets using scRNAseq analyses, and 85 genes were obtained by overlapping the OSM target genes from NicheNet with genes significantly highly expressed in ICC cells (Figure 5A). Among these 85 genes, pathways related to response to cytokine, MAPK cascade, epithelial cell development, and cell proliferation were enriched (Figure 5B). Enrichment results of differentially phosphorylated proteins further validated the enrichment of MAPK and cell cycle pathways in exogenous OSM-treated QBC939 cells (Figure 5C). Subsequent protein interaction analysis revealed that Osmr interacted with Stat3, and Stat3 interacted with Ccnd1 (Figure 5D). IHC results revealed that the expression levels of p-STAT3 and CCND1 were significantly decreased in Osmr-knockout mouse ICC tissues (Figure 5E). Similarly, Western blotting results indicated Osmr knockout decreased CCND1, BCL2, and p-STAT3 protein levels (Figure 5F). Therefore, OSM appears to promote ICC growth via OSMR and downstream CCND1, BCL2, and p-STAT3 signaling.

**Figure 5.**
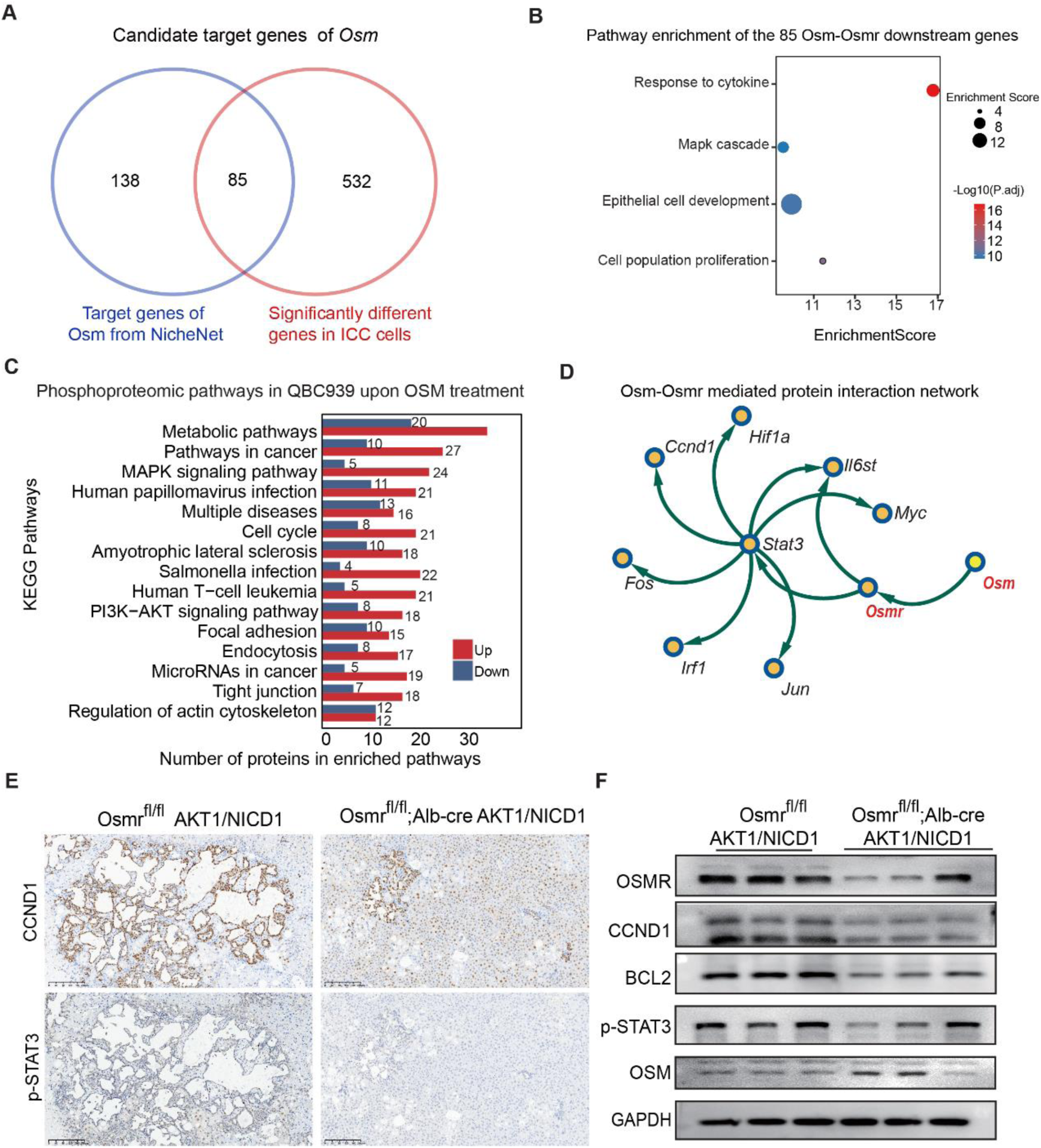
Mφ_VEGFA promotes the proliferation of ICC cells by activating p-STAT3 and CCND1/MYC signaling via OSM-OSMR. **(A)** A Venn plot displays the candidate target genes of Osm. **(B)** Enriched pathway analysis of the candidate target genes. **(C)** Protein network demonstrating the candidate downstream signaling of Osm-Osmr. **(D)** Immunohistochemical staining of p-STAT3 in paraffin-embedded liver sections from Osmr^fl/fl^ and Osmr^fl/fl^; Alb-Cre mice injected with AKT1/NICD1. Scale bars represent 50 μm. **(E)** Immunohistochemical staining of CCND1 and p-STAT3 in liver tissues from Osmr^fl/fl^ and Osmr^fl/fl^; Alb-Cre mice injected with AKT1/NICD1. Scale bars, 200 μm. **(F)** Western blot analysis of OSMR, CCND1, BCL2, phosphorylated STAT3 (p-STAT3), and OSM protein levels in Osmr^fl/fl^ and Osmr^fl/fl^; Alb-Cre mice injected with AKT1/NICD1.

### Mφ_VEGFA promotes ICC invasion via THBS1*–*CD47*–*mTOR*/*DBN1 axis

Next, we investigated the roles of THBS1 in ICC progression and found that THBS1 interacts with ICC cells via CD47 (Figure 6A, Supplementary Figure S9A–B). Using NicheNet, we identified PI3K–AKT and EMT-related genes as potential *THBS1* downstream signaling in ICC cells (Figure 6A). Critically, recombinant THBS1 treatment marginally favored ICC cell growth (Supplementary Figure S9C–D), but markedly enhanced ICC cell migration (Figure 6B–C). Correspondingly, silencing CD47 significantly impaired the stimulatory effects of THBS1 on ICC cell migration (Figure 6D). Co-culture with BMDMs increased BR migratory ability, whereas *Cd47* knockdown blocked this effect (Figure 6E–F, Supplementary Figure S9E). *Cd47* silencing inhibited subcutaneous allograft progression in mice (Figure 6G, Supplementary S9F–G). Moreover, CD47 knockdown markedly suppressed ICC dissemination in a peritoneal metastasis mouse model, as evidenced by reduced metastatic nodule formation and decreased ascites accumulation (Figure 6H, Supplementary Figure S9H). Consistently, CD47 silencing attenuated ICC growth and local invasion into the surrounding hepatic parenchyma in the orthotopic allograft ICC mouse model (Figure 6I). These results together indicate that THBS1 promotes ICC cell migration via its receptor CD47.

**Figure 6.**
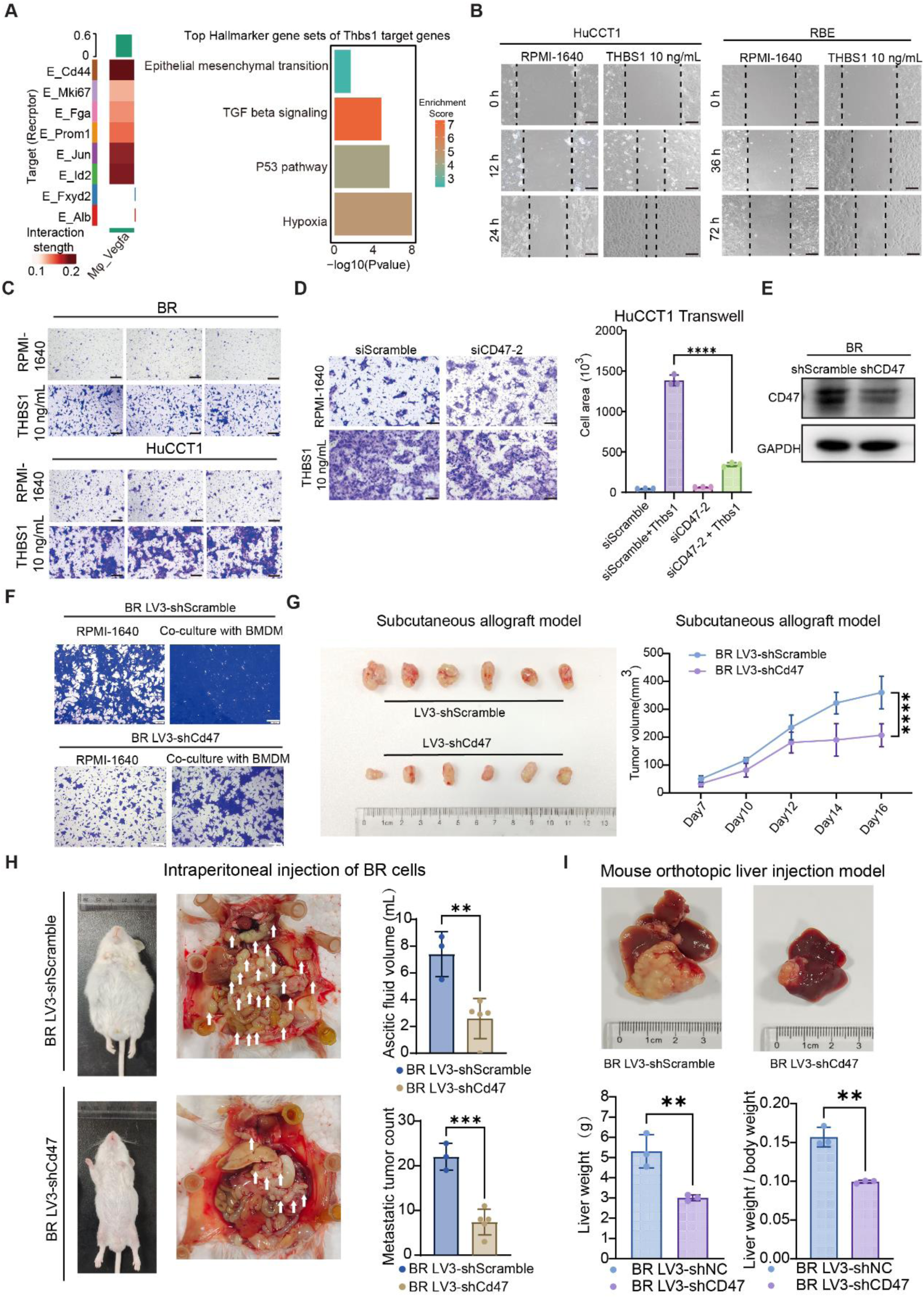
Mφ_VEGFA promotes ICC invasion via THBS1–CD47 signaling. **(A)** Heatmap of interactions between Mφ_Vegfa and normal and malignant epithelial subtypes via THBS1 signaling (left), and bar plot demonstrating the hallmark target gene sets of *Thbs1* obtained by NicheNet (right). **(B)** Wound healing assay of HuCCT1 and RBE cells treated with or without 10 ng/mL THBS1. HuCCT1 cells collected at 0, 12, and 24 h; RBE cells collected at 0, 36, and 72 h. Scale bar, 150 μm. **(C)** Transwell migration assay of BR (mouse) and HuCCT1 (human) cells treated with 10 ng/mL THBS1. Scale bar, 150 μm. **(D)** Transwell migration assay of HuCCT1 cells transfected with siCD47-2 and treated with or without 20 ng/mL THBS1 for 48 h.**(E)** Western blot of CD47 in BR cells infected with LV3-shNC or LV3-shCD47 lentivirus. **(F)** Transwell migration images of BR cells infected with LV3-shNC or LV3-shCd47 lentivirus and co-cultured with BMDMs for 24 h. Scale bar, 150 μm. **(G)** Images of subcutaneous allografts and allograft growth curves of BR cells infected with LV3-shNC or LV3-shCd47 lentivirus. **(H)** Representative gross images of mice with intraperitoneal injection of LV3-shNC (n = 3 mice) or LV3-shCd47 infected BR cells (n = 5 mice). **(I)** Representative images of mouse abdominal ICC nodules at the end of the experiment after intraperitoneal injection of BR cells infected with LV3-shNC or LV3-shCd47. Tumor nodules were indicated by white arrows. Data are presented as mean ± SD. Significance was determined by the two-tailed unpaired Student t-test for two groups or by one-way ANOVA followed by Dunnett’s t-test for multiple groups. ** *P* < 0.01, *** *P* < 0.001, and **** *P* < 0.0001.

To further elucidate the downstream molecules of *THBS1/CD47* in promoting ICC migration, we treated HuCCT1 cells with THBS1 and collected cells at different time points. Western blotting demonstrated that treatment with THBS1 induced rapid induction of EMT-related proteins N-Cadherin and Vimentin around 2 hours (Figure 7A, Supplementary Figure S10A). Phosphoproteomics analysis of HuCCT1 cells treated with recombinant THBS1 at 0 and 2 h revealed elevated phosphorylation of PI3K–AKT and EMT-related proteins at 2 h (Figure 7B). Treatment of ICC with THBS1 showed that p-mTORC1 and p-RPS6 were significantly increased (Figure 7C, Supplementary Figure S10A). mTORC1 inhibition by rapamycin substantially suppressed the pro-migratory effect of BMDM co-culture on ICC cells and completely abrogated the THBS1-induced ICC cell migration (Figure 7D–E), indicating that THBS1 promotes ICC cell migration via mTORC1 activation.

**Figure 7.**
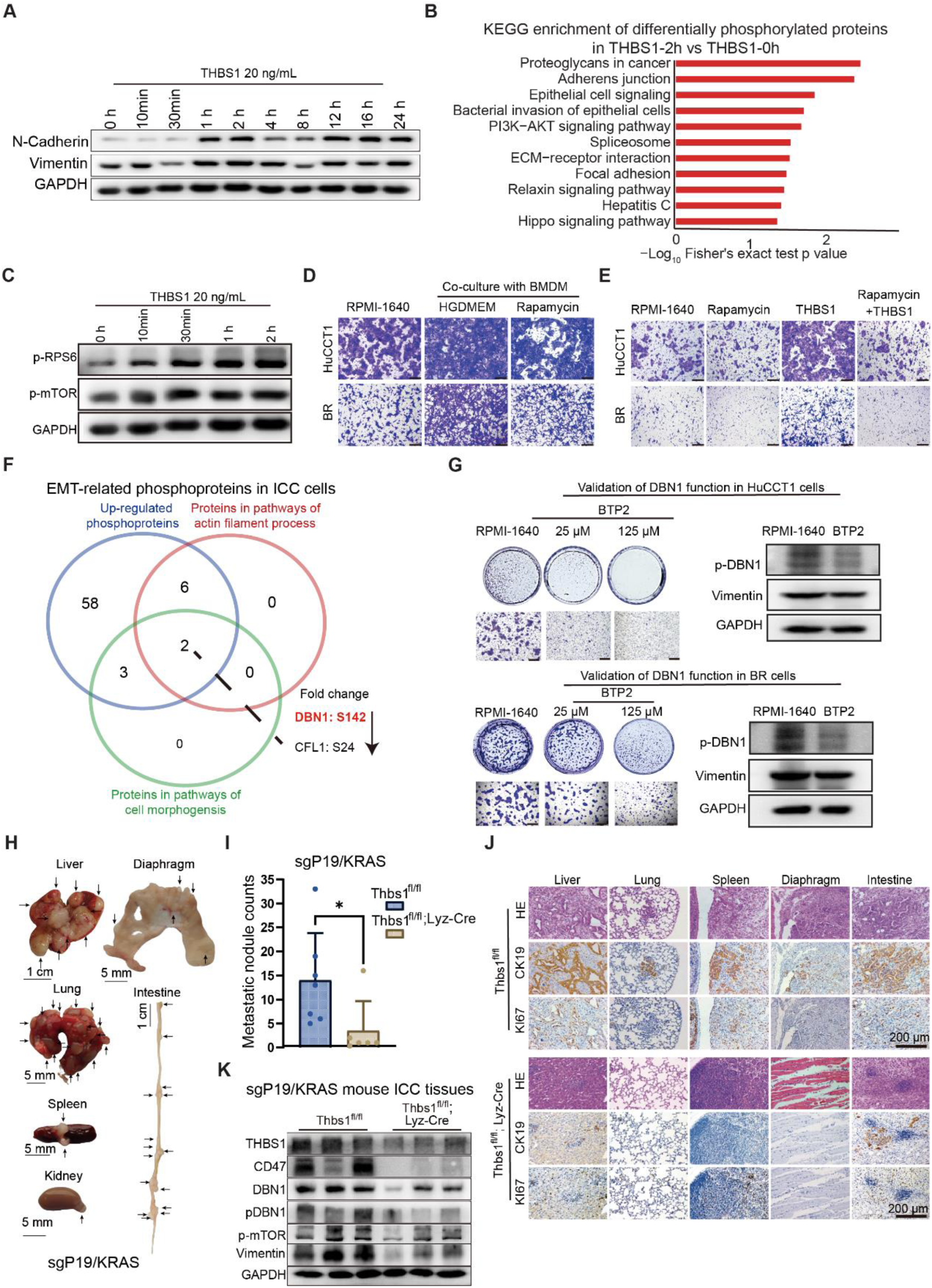
Mφ_VEGFA derived THBS1 promotes ICC invasion via activated mTOR and phosphorylated DBN1. **(A)** Western blot analysis of N-cadherin and Vimentin expression in HuCCT1 cells treated with 20 ng/mL THBS1 at the indicated time points. GAPDH served as a loading control. **(B)** Bar plot displaying pathway enrichment analysis for HuCCT1 at 2 h following addition of exogenous THBS1. **(C)** Western blot of p-RPS6 and p-mTOR in BR cells cultured alone or co-cultured with BMDMs at the indicated time points. GAPDH served as a loading control. **(D)** Transwell migration assays of HuCCT1 and BR cells co-cultured with BMDMs and with or without rapamycin treatment. N = 5. Scale bar, 200 μm. **(E)** Transwell migration assays of HuCCT1 and BR cells treated with RPMI-1640, rapamycin, THBS1, or THBS1 + rapamycin. Scale bar, 200 μm. **(F)** Venn plot showing the enriched phosphorylated proteins involved in the EMT process. **(G)** Transwell migration assays for HuCCT1 and BR cells treated with RPMI-1640, 25, or 125 μM BTP. Scale bar, 200 μm. Western blot analysis of p-DBN1 (Ser142) and Vimentin in HuCCT1 and RBE cells treated with 50 μM BTP2 for 24 h. GAPDH was used as a loading control. **(H)** Representative images of metastatic ICC lesions from the indicated organs in Thbs1^fl/fl^; Lyz-Cre and Thbs1^fl/fl^ mice injected with sgP19/KRAS. **(I)** Metastatic nodule numbers in the sgP19/KRAS ICC metastasis model. Data are presented as mean ± SD. Significance was determined by a two-tailed unpaired Student’s t-test (\**P* < 0.05). **(J)** H&E, CK19, and Ki67 staining of metastatic tumor nests in sections from different organs in Thbs1^fl/fl^ mice and Thbs1^fl/fl^; Lyz-Cre mice injected with sgP19/KRAS. This lymph node is the largest adjacent to the spleen. **(K)** Western blot analysis of THBS1, CD47, DBN1, p-DBN1(Ser142), p-mTORC1(Ser2448) and Vimentin protein levels in ICC tissues from Thbs1^fl/fl^; Lyz-Cre and Thbs1^fl/fl^ mice injected with sgP19/KRAS. GAPDH served as a loading control.

In addition to m-TORC1, DBN1 was identified as the first phosphorylated protein by fold change in cytoskeletal remodeling and EMT-related signaling pathways via phosphoproteinomics, and Ser142 as the most strongly upregulated phosphorylation site (Figure 7F). Likewise, DBN1 inhibition with its inhibitor BTP2 significantly blocked ICC cell migration (Figure 7G). Notably, BTP2 decreased phosphorylation at Ser142 of DBN1 (Figure 7G). Collectively, these results demonstrate the critical roles of p-mTORC1 and p-DBN1(Ser142) in favoring THBS1-mediated ICC cell migration.

Finally, we examined the role of THBS1 in the sgP19/KRAS-induced metastatic ICC mouse model with Thbs1^fl/fl^; Lyz-Cre mice (Figure 7G). Myeloid-specific knockout of *Thbs1* significantly suppressed peripheral metastasis of ICC to multiple distant organs, including the livers, lungs, spleens, kidneys, diaphragm, and intestines (Figure 7H–I). H&E and IHC staining confirmed the metastasis of ICC in those organs (Figure 7J). Furthermore, Western blotting of ICC tissues revealed that metastasis-related CD47, DBN1, p-DBN1 (Ser142), p-mTORC1, and Vimentin levels were significantly downregulated after *Thbs1* deletion (Figure 7K). Therefore, THBS1 appears to promote ICC metastasis via PI3K-AKT-mTOR activation and DBN1 phosphorylation.

### Blockade of OSM and knockout of THBS1 synergistically inhibit ICC progression

As *OSM* promotes ICC cell proliferation and *THBS1* enhances invasiveness, we were curious whether dual inhibition could synergistically suppress ICC development. To this end, we first analyzed synergistic expression of *THBS1* and *OSM* in 827 human ICC tissues across nine ICC/cholangiocarcinoma cohorts by calculating mean expression levels of *THBS1* and *OSM* in each cohort (Supplementary Table S2). An ICC sample was defined as double-positive if both genes had expression levels above the mean of the whole cohort. Across the nine datasets, the proportion of ICC samples co-expressing *THBS1* and *OSM* ranged from 13.33% to 29.53%, and those in five datasets exceeded 20% (Figure 8A). Prognostic analyses across three datasets demonstrated that patients with high expression of both *OSM* and *THBS1* had significantly shorter overall survival (Figure 8B). mIF staining indicate that OSM^+^THBS1^+^CD68^+^VEGFA^+^ cells were spatially adjacent to CK19^+^ cells (Figure 8C), and OSM^+^THBS1^+^ were co-expressed in CD68^+^ cells (Figure 8D). Correspondingly, patients of ICC with highly co-expression of OSM^+^THBS1^+^ exhibited significantly worse survival, indicating that dual expression of OSM and THBS1, widely present in ICC, was closely related to the outcome of patients with ICC (Figure 8E).

**Figure 8.**
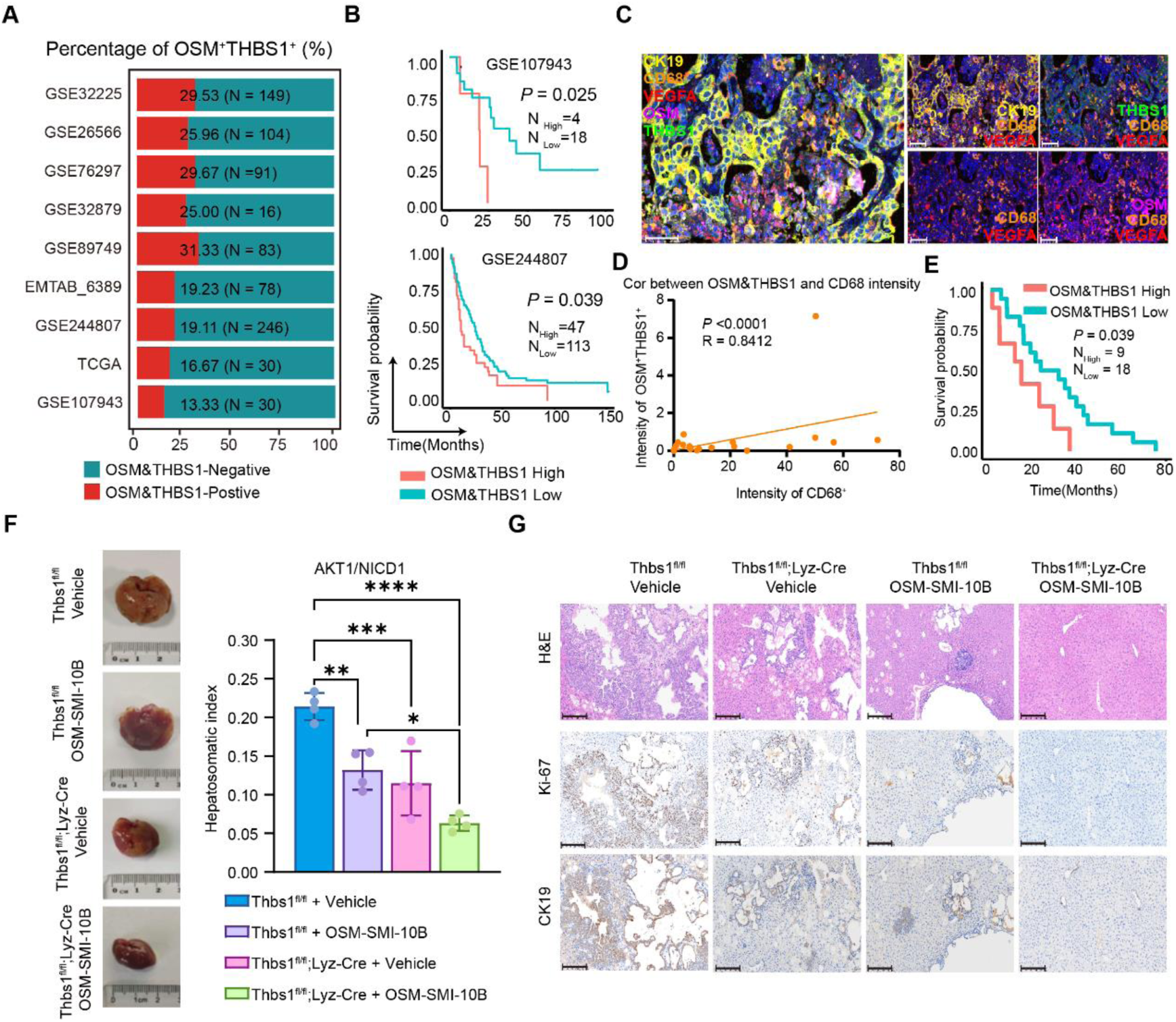
Combined intervention with OSM inhibitor and THBS1 knockout delays ICC progression. **(A)** Stacked bar plot illustrating the prevalence of OSM⁺THBS1⁺ dual-positive ICC within distinct cohorts. **(B)** Survival curve of patients with ICC with high or low OSM and THBS1, respectively. **(C)** mIF shows OSM^+^THBS1^+^CD68^+^VEGFA^+^ macrophages around CK19^+^ cells. Scale bar, 50 μm. **(D)** Spearman correlation between OSM^+^THBS1^+^ and CD68^+^ intensity. **(E)** Patients were stratified by OSM^+^ and THBS1^+^ intensity (red = OSM^+^THBS1^+^ high, green = OSM^+^THBS1^+^ low); survival was compared using the log-rank test. **(F)** Representative liver images from mice with THBS1 knockout, OSM inhibition, or combination blockade as indicated. Statistical analysis of liver-to-body weight ratio across different treatment groups in the *THBS1/OSM* combination blockade experiment, n = 4 mice. Data are presented as mean ± SD. Statistical significance was analyzed using one-way ANOVA (*p* < 0.05), followed by Dunnett’s t-test (*\*P* < 0.05, *\*\* P* < 0.01, *\*\*\* P* < 0.001; *\*\*\* P* < 0.0001). **(G)** H&E, Ki67, and CK19 staining of liver sections from the same mice as stated in panel F. Scale bar, 200 μm.

Therefore, we genetically ablated macrophage-derived *THBS1* using Thbs1^fl/fl^; Lyz-cre mice and pharmacologically inhibited OSM-OSMR interaction via intraperitoneal administration of OSMR inhibitor OSM-SMI-10B to examine their synergistic effect (24). Intriguingly, *Thbs1* deletion or *Osmr* inhibition alone suppressed ICC progression moderately, while combined blockade attenuated ICC development more robustly, as evidenced by further reduced tumor burden and decreased liver-to-body weight ratio (Figure 8F–G). Therefore, macrophage-derived OSM and THBS1 may represent promising therapeutic targets, and combined blockade of OSM and THBS1 may yield synergistic therapeutic effects in a substantial subset of patients with ICC.

## Discussion

TAMs in ICC have been associated with poor prognosis through tumor growth and invasiveness promotion(25, 26). Recent single-cell omics research has revealed the molecular diversity and functions of TAMs in cancer (27). Nonetheless, the specific subgroups driving ICC malignancy and its interactions with tumor cells remain unclear. Using a time-resolved ICC mouse model with integrated multi-omics, we identified Mφ_VEGFA as a poor-prognosis TAM subset in ICC. Notably, the prognostic significance of Mφ_VEGFA was also observed in hepatocellular carcinoma (13), underscoring its prevalence across liver cancers (28).

Moreover, our study uncovers the mechanism underlying Mφ_VEGFA to promote ICC progression or metastasis instead of suppressing the tumor immune microenvironment. Specifically, Mφ_VEGFA promotes ICC cell proliferation through OSM-OSMR and downstream p-STAT3 and CCND1, corroborating the findings that OSM promotes cancer cell proliferation in glioblastoma (29) and pancreatic ductal adenocarcinoma (30). Conversely, Mφ_VEGFA exerts its pro-invasive role through secreted THBS1 to activate CD47 signaling in ICC cells. Notably, THBS1 has been reported to be released from tumor-infiltrating monocyte-like cells and to induce an immunosuppressive TME to boost metastasis (31). Conversely, we demonstrate that *CD47* activates mTOR and DBN1 in ICC cells but not IL6/STAT3, causing EMT and cell skeleton remodeling (32).

Finally, our findings offer a new therapeutic strategy for ICC treatment by targeting molecules derived from Mφ_VEGFA, which secrete OSM and THBS1 to accelerate ICC growth and metastasis via OSMR and CD47, respectively. Combined blockades of OSM–OSMR and THBS1–CD47 signaling retarded growth and restrained metastasis in primary ICC mouse models, while single inhibition generated modest efficacy. Consistently, a clinical trial demonstrated that targeting OSM-OSMR signaling alone led to poor outcomes or drug resistance (33). Additionally, OSM and THBS1 were concurrently upregulated in 13–29% of patients with ICC. Thus, targeting dual modulators of Mφ_VEGFA provides a potentially more robust therapeutic option for those patients with ICC.

Nonetheless, this study also has some limitations. First, the role of Mφ_VEGFA in promoting ICC migration and invasion was not fully illustrated using matched early and late-stage clinical samples. As ICC is usually diagnosed at advanced stages and progresses rapidly, it is difficult to obtain paired pre- and post-metastatic ICC tissues from the same patients. Therefore, it is challenging for us to determine the prevalence of Mφ_VEGFA in metastatic ICC. Second, we were unable to explore the possibility of specific Mφ_VEGFA subpopulation depletion to confirm its unique role in favoring ICC progression. Clearly, this caveat is caused by a technical limitation, as no reliable technique is available to delete a specific macrophage subpopulation to date. Finally, the therapeutic value of targeting OSM or THBS1 was not fully demonstrated in this study. In this regard, organoid or patient-derived xenograft mouse models are more suitable models for assessing the therapeutic significance of specific targets. However, these models are not yet well established in our laboratory. Therefore, future studies should consider using these models to overcome these limitations and further evaluate the feasibility of targeting OSM/THBS1 for ICC treatment.

## Methods

### Sex as a biological variable

Only female mice were used in this study, and sex was not considered a biological variable for study design and data analysis.

### Animals

Wild-type FVB and C57BL/6 mice were purchased from Beijing Vital River Laboratory Animal Technology Co., Ltd. (Beijing, China). Genetically modified mice were provided by Shulaibao Biotechnology Co., Ltd. (Wuhan, China). All mice were maintained in a specific pathogen-free environment. Detailed information regarding the animals used in this study is provided in Supplementary Table S1.

### Cell source

The human cholangiocarcinoma cell lines HuCCT1, RBE, and QBC939, as well as the mouse fibroblast cell line L929 and macrophage cell line RAW 264.7, were used in this study. HuCCT1 was obtained from the Japanese Collection of Research Bioresources Cell Bank (JCRB, Osaka, Japan; accession number JCRB0425). RBE was acquired from the China Center for Type Culture Collection (Wuhan, China; accession number GDC0765). QBC939 was purchased from Shanghai Jinyuan Biological Technology Co., Ltd. (Shanghai, China; product code JY-Y1457). The mouse cell lines L929 (ATCC® CCL-1™) and RAW 264.7 (ATCC® TIB-71™) were purchased directly from the American Type Culture Collection (ATCC, Manassas, VA, USA). All cells were cultured according to the providers’ recommendations and were routinely tested for mycoplasma contamination.

BR cells, isolated from BMI1/NRAS–driven primary ICC tissues in FVB mice, were maintained in RPMI-1640 complete medium supplemented with 10% fetal bovine serum and 1% penicillin-streptomycin at 37°C with 5% CO₂ in an incubator. These cells displayed consistent *in vitro* growth characteristics and were capable of forming tumors in both subcutaneous and orthotopic liver implantation models in wild-type FVB mice. IHC and histopathological analysis confirmed the presence of ICC features.

### Establishment of ICC Mouse Models

Primary ICC mouse models were established in 6–8-week-old mice via hydrodynamic tail vein injection. For the AKT1/NICD1 ICC model (NICD1, Notch1 intracellular domain), a plasmid mixture containing 15 μg pT3-EF1aH-myr-AKT, 15 μg pT3-EF1a-NICD1, and 1.5 μg pCMV-SB, was diluted in 2 mL of sterile saline and filtered through a 0.22 μm membrane to remove impurities. For the highly metastatic sgP19/KRAS ICC model (34), a mixture of plasmids, including 10 μg CRISPR-Cas9-sgP19, 25 μg pT3-EF1α-KRAS, and 5.0 μg pCMV-SB, was diluted in 2 mL of sterile saline. For a 20 g mouse, the injection volume was 2 mL and was adjusted according to body weight. Mice were restrained using a tail vein visualization platform, and the injection was completed within 5–7 s. Mice were euthanized and dissected either at the end of the experiment or when moribund.

### Public single-cell and spatial transcriptomics data collection

In addition to in-house single-cell and spatial transcriptomics data, we also curated a an AKT1/YAP1-driven ICC mouse model(15), two human ICC cohorts single cell transcriptomic data (GSE138709(16) and GSE189903(17)), and a human spatial transcriptome data using 10x Visium platform (23) to validate our finding in the AKT1/NICD1 mouse ICC tissues (Supplementary Table S2).

### Single-cell transcriptome data processing and cell type definition

For in-house AKT1/NICD1, public AKT1/YAP1, and human ICC single cell transcriptome data, gene expression matrices were preprocessed using the Seurat R package (V 4.4.0)(35). Quality control measures were employed to remove low-quality cells and genes based on the following criteria: 1) remove ribosomal genes. 2) nCount_RNA > 1000, nCount_RNA < 40000, nFeature_RNA < 6000, nFeature_RNA > 500, percent.mt < 20%, min.cells=50. DoubletFinder(36) (V 2.0.6, default parameters) to remove doublet cells. Then, the global-scaling method “LogNormalize” normalized the gene expression measurements for each cell by the total expression. The ScaleData function, “vars.to.regress” option percent mitochondrial content was used to regress out unwanted sources of variation. Top highly variable genes (HVGs, vst, top 2000) were calculated using the Seurat function FindVariableGenes. Principal component analysis (PCA) for dimensionality reduction was performed using the top 30 components. Clusters were computed using the FindClusters function using the top 30 PCAs and then visualized using the uniform manifold approximation and projection (UMAP) implemented in Seurat. Differential expression analysis between clusters was performed using the Wilcoxon rank sum test (RunPrestoAll in the presto R software) at a family-wise error rate of 5%.

Using canonical marker genes (Supplementary Table S3), SciBet (V1.0) (37) and differentially expressed genes (DEGs) and the associated pathway function to define the major cell types (Fibroblasts, Endothelial, Macrophage, DC, Neutrophile, T&NK, B, Cholangiocarcinoma, and Hepatocyte). Sub-cell types were re-clustered with a resolution parameter set at 0.2–0.5, without filtering cells or genes, and the data were processed using the standard workflow to identify the subclusters and their associated differential genes (HVGs, vst,1000 for human macrophage and epithelial cells, 500 for AKT1/NICD1 epithelial cells, 300 for AKT1/NICD1 and AKT1/YAP1 mouse macrophage).

In each subtype, if the main cell type marker genes from the first step are simultaneously expressed at a high proportion (>30% of cells) and with high specificity in a cell subpopulation, then that subpopulation is removed. Meanwhile, we used SciBet to remove mixed cells to ensure the accuracy of the final identified cell types. Scibet references downloads from website http://scibet.cancer-pku.cn/scibet_references.html (human:30 major human cell types, mouse: Single-cell transcriptomics of 20 mouse organs creates a Tabula Muris). Finally, based on the functions of the differentially expressed genes and the cell type marker genes, subgroups with similar functions were merged, and the cell types were defined.

For human ICC single cell transcriptome data from GSE138709 and GSE189903, we employed the same quality control, normalization, and dim reduction as AKT1/NICD1 and AKT1/YAP1. Additionally, we merged the two cohorts and used Harmony (38) (V1.1.0) to eliminate batch effects caused by different platforms and sequencing protocols. Cell type definition was the same as in the AKT1/NICD1 and AKT1/YAP1 data.

### Enrichment score of each cell type

To assess the relative enrichment of cell subpopulations across different stages of ICC progression, we calculated the ratio of observed to expected cell numbers (Ro/e) for each cell subset (39). The expected number of cells for each subset was determined using χ² tests. Ro/e > 1 indicated enrichment of a given cell subset at a particular stage of ICC progression, suggesting a potential association with that disease stage.

### Macrophage subtype-associated clinical prognostic integrative analysis

To predict the association between macrophage subtypes and survival phenotypes of ICC patients, we utilized the R package Scissor (v2.0.0) (18) to link single-cell data with patient phenotypes from bulk sequencing datasets GSE89749 (19) and GSE107943 (20). In detail, we analyzed the correlation between macrophage subtypes and the prognostic data of ICC patients via Scissor tutorial, employed Cox-regression, and selected overall survival as the dependent variable. For each macrophage cell type, we set parameter alpha 0.2 and calculated the fractions of Scissor+ cells (positively correlated with worse survival), Scissor– cells (positively correlated with favorable survival), and neutral cells (uncorrelated with survival).

Meanwhile, we validated the Mφ_VEGFA expression signature in GSE89749 and GSE107943 cohorts. The top 10 differentially upregulated marker genes of Mφ_VEGFA (ranked by Log2FC) were selected (Supplementary Table S4). A Mφ_VEGFA score was calculated as the mean expression of these genes per patient. Patients were stratified into Mφ_VEGFA-High and Mφ_VEGFA-Low groups based on the median score. The association between the Mφ_VEGFA score and overall survival (OS) was then evaluated. Statistical significance was determined using the log-rank test (Mantel-Cox), with P values < 0.05 considered statistically significant.

### Cell type communication analysis

We conducted a comprehensive computational analysis of cell-cell communication to decipher the ligand-receptor interactions between macrophage and tumor subtypes, utilizing the bioinformatics tools CellphoneDB (40), CellChat (41), and NicheNet (42). By integrating the results from these three methods, we systematically identified key communication components, downstream signaling pathways, and target genes involved in the interactions between macrophages and tumor cells.

We utilized CellChat to analyze the ligand-receptor gene pairs and signaling pathways involved in communication between macrophage and tumor cell subgroups. Cell communication pairs with significant p-values (*P* < 0.05 and prob > 0.01) were identified as candidate gene pairs. In the CellPhoneDB package (V3), we selected receptors and ligands expressed in over 10% of cells in the specific cluster for subsequent analysis. In mouse single-cell data, we obtained the corresponding human orthologs using the convert_mouse_to_human_symbols function in the nichenetr R package (V2.1.0). We selected the interactive pairs with interaction mean scores > 1.0 and *p* < 0.05.

Furthermore, we used NicheNet to analyze interactions between macrophage and tumor cell subgroups, as well as the potential downstream target genes regulated by ligand genes. In this pipeline, we selected genes expressed in at least 10% of the cells as the background gene set, selected differential expression genes in macrophage and tumor subtypes (log2FC > 0.3 and p.adj < 0.05) as the gene set of interest. NicheNet’s ligand-receptor network and ligand-target matrix were used to predict active ligand-receptor pairs and rank ligands by their regulatory potential for macrophage and tumor subtypes. We obtained active ligand–receptor gene pairs using the predict_ligand_activities function. Parametric Pearson correlation analysis was performed to assess the correlation between each ligand and its downstream pathway target genes, and ligands with a Pearson correlation coefficient above the lower quartile were retained as active ligands. Ligand-associated downstream target genes were subsequently extracted from the built-in NicheNet database.

The final selection of candidate ligands was based on those identified by all three software tools, with an additional requirement that their expression levels in human and mouse Mφ_VEGFA subtype were significantly high (Log_2_FC > 0.3).

### CNV analysis

CNV (copy number variation) scores were estimated in every single cell based on averaged expression profiles across chromosomal intervals using infercnv (43) R package (V1.0.4). The CNV score was calculated as the quadratic sum of the CNV region (44).

### Spatial transcriptomics analysis

#### Data processing

For ICC AKT1/NICD1 Day 31 mouse spatial transcriptomics data, we performed cell segmentation and obtained spatial gene expression matrices via BSTMatrix (v2.3, combined spot level=7) using the GRCm39 genome reference.

Next, for human ICC tissue using 10x Visium and AKT1/NICD1 mouse ICC spatial transcriptome data, we processed the expression matrix via Seurat (V4.4.0), and retained spots with over 200 genes, mitochondrial content below 20%, and haemocyte below 5%. Then, we normalized the count matrices by sctransform method. Next, we applied the top 30 PCAs and obtained spatial clusters. Subsequently, we identified DEGs by the FindAllMarkers function and performed functional enrichment analysis with clusterProfiler (45) (V4.10).

For in-house human ICC spatial transcriptomic data generated on the 10x Visium HD platform, the raw expression matrix was obtained from square_016um bins. Data processing was performed using the sketch-based workflow in Seurat (v5.3.1). Spots with fewer than 20 detected genes, mitochondrial content above 20%, or haemocyte content above 5% were excluded. The top 50 principal components were used for clustering. Differentially expressed genes for each cluster were identified using the FindAllMarkers function in the Seurat R package.

### Cell type defined

For high-resolution spatial data using BMKMANU S1000 AKT1/NICD1 and 10x HD platform, we employed spacexr (46) (V2.2.1) to integrate scRNAseq data to define the cell types of each cluster. For BMKMANU S1000 AKT1/NICD1 spatial data, we integrated scRNAseq data from Day 31 AKT1/NICD1 mouse tissues using the “full” model in spacexr. For in-house 10x Visium HD spatial transcriptomic data from four human ICC tumor tissues, scRNA-seq references were compiled by merging GSE138709 and GSE189903, retaining only cell types enriched in tumor tissues. Neutrophils were excluded from downstream integration analysis, as they were present at very low abundance in the FFPE ICC samples. Cell type deconvolution across spatial spots was then performed by integrating the spatial data with the scRNA-seq reference using the doublet model in spacexr.

### Cell type scores

Cell type scores for individual Mφ_VEGFA and epithelial subtypes in spatial transcript data were evaluated via Seurat’s AddModuleScore function (ctrl = 5) using the cell type feature genes listed in the Supplementary Table S4. The resulting enrichment scores were subsequently visualized utilizing the standard FeaturePlot and VlnPlot functions.

### Spatial segment

Spatial spots were first classified as “Tumor” or “Normal” by inferring somatic CNVs using InferCNV. Spots designated as normal epithelial or stromal references were used to define the baseline CNV state. Clusters exhibiting large-scale chromosomal gains or losses relative to the reference were annotated as “Tumor”, while clusters with flat CNV profiles were annotated as “Normal”. Clusters that could not be confidently assigned to either category were excluded from subsequent boundary analysis.

To delineate the tumor – normal boundary, boundary-proximal spots were first identified using k-nearest neighbor (KNN) analysis (k = 30) on pixel coordinates: a spot was designated as a boundary spot if it belonged to one category while at least one of its k neighbors belonged to the opposing category.

A continuous probability field was then estimated across the tissue section. For each point on a 500 × 500 grid spanning the full coordinate space, KNN regression (k = 30) was applied with binary Tumor/Normal labels (1/0) as the response variable. The resulting probability surface was smoothed with a Gaussian kernel via the isoblur function from the imager package. Grid regions farther than 300 pixels from any sequenced spot were masked to restrict the boundary to tissue-covered areas. The tumor boundary was extracted as the 0.5-probability isoline using contourLines, retaining only contour segments with ≥ 150 vertices; the longest valid contour was selected as the final boundary polyline.

Each spot was assigned a perpendicular distance to the boundary polyline via point-to-segment projection. To account for the spatial resolution of each platform, multiple concentric bands were defined on both sides of the boundary: 500 μm for human ICC tissue profiled on the 10x Visium platform, and 100 μm for AKT1/NICD1 mouse ICC tissue profiled on the BMKMANU S1000 platform. Spots falling within these bands were collectively designated as the “Tumor Boundary” zone, while spots outside this region retained their original Tumor or Normal annotation, yielding three tissue segments—Tumor, Tumor Boundary, and Normal— for downstream analysis.

### BANKSY analysis

For 10x HD spatial transcription data, we conducted spatial-domain analysis using BANKSY (21) in the Seurat 5.3.1 package. Specifically, we utilized the expression matrix of each sample, post-preprocessing, along with the spatial coordinate information of the spots as input. BANKSY was then applied with default resolution parameters to assign each spot to spatially informed clusters across all samples.

### MistyR analysis

After cell type definition, we employed mistyR (22) to estimate the importance of the abundance of each cell type in explaining the abundance of other cell types by extracting relationships from spatial transcriptome data through multiple views focusing on different spatial or functional contexts. We defined an intraview, local niche view (juxtaview), where the radius is the mean distance to the nearest neighbor plus one standard deviation, and a broader tissue view (paraview), where the radius is the mean distance to the 7th nearest neighbor plus one standard deviation, to capture the distribution of cell type proportions within the neighboring tissue. To identify cell type communities, we processed the “importances. aggregated” output from mistyR, which consists of aggregated, view-specific predictor-target importance tables. We then filtered the Paraview “importances.aggregated” using a threshold of 0.5, setting values below this threshold or NA values to zero.

### Statistics

All experiments were conducted with rigorous reproducibility, including at least three independent biological replicates per group (n ≥ 3) to ensure result stability and reliability. Technical replicates were performed for each sample to minimize experimental error. For animal studies, each group included no fewer than three animals (n ≥ 3), all treated and sampled under identical conditions.

Statistical analyses were performed using GraphPad Prism. Comparisons between two groups were conducted using two-tailed unpaired Student’s t-tests. For statistical comparison of more than two groups, one-way ANOVA with the Tukey post hoc test was used. Significance levels are indicated as follows: \**P* < 0.05, \*\**P* < 0.01, \*\*\**P* < 0.001, \*\*\*\**P* < 0.0001, and ns: not significant. All data are representative of three biologically independent experiments, unless otherwise indicated.

## Study approval

This study was conducted in accordance with the Declaration of Helsinki and the Declaration of Istanbul. All procedures involving animals were approved by the Animal Experiments Ethics Committee of Huazhong University of Science and Technology (approval No. 20230312). Tissue array was purchased from Shanghai Outdo Biotech (Shanghai, China) with ethical approval No. SHYJS-BC-2111001-06 (GZ). Tissue array for THBS1^+^ clinical analysis was purchased from the Shanghai LiaoDing Biotechnology (Shanghai, China) with ethical approval No. 2020-035-01. All patients provided written informed consent before sample collection.

## Data Availability

The scRNA-seq data generated in this study are publicly available in the GEO database under accession code GSE308574. The remaining data are available within the manuscript, supplementary information, or from the corresponding author upon reasonable request.

## Other Methods

Methodological details for single-cell RNA-seq, single-cell ATAC-seq, spatial transcriptomics, proteomics, and phosphoproteomics, as well as all in vivo and in vitro experiments, are provided in the Supplementary Materials and Methods.

## Authors’ Contributions

C.X. and T.W. designed the study. T.W. analyzed the multi-omics data. R.L. performed the experiments. T.W. and R.L. wrote the manuscript. C.X. revised the manuscript. Z.C. and M.Z. prepared the samples. B.W. and C.L. managed data collection and organization. All authors have read and approved the manuscript.

## Funding support

This research was supported by grants from the National Natural Science Foundation of China (82472988, 82273059, and 82573814), the Henan Province and Ministry of Health of Medical Science and Technology Program (SBGJ202302028 for T.W. and SBGJ202402032 for C.L.), and the Fundamental Research Funds for the Central Universities (2025JYCXJJ067 for M.Z.).

## Acknowledgments

This work was financially supported by the National Natural Science Foundation of China (Grant Nos. 82472988, 82273059, and 82573814).

## Supplemental material

To access the supplementary material accompanying this article, visit the online version.

## Conflict of Interest Statement

The authors have declared that no conflict of interest exists.

## Notes

### Competing Interest Statement

The authors have declared no competing interest.

### Summary of Updates

The conclusions have been updated to better reflect the findings, and all main figures have been revised and optimized.

