## Supplementary material for "VEGFA+ macrophages promote the growth and metastasis of intrahepatic cholangiocarcinoma via OSM and THBS1 signaling": sup.pdf

### Supplementary Materials and Methods

#### Supplementary Methods

##### Single-cell RNA sequencing

Under sterile conditions, the freshly collected liver was washed twice with pre-cooled RPMI-1640 medium containing 0.04% BSA. The tissues were then mechanically dissociated into approximately 0.5 mm³ fragments using surgical scissors and transferred into a freshly prepared enzymatic digestion solution. The digestion mixture contained RPMI1640 (Conring, cat. no. 10-040-CVR), 0.04% BSA (MACS, cat. no. 1000076), and 0.2% collagenase II (Gibco, cat. no. 17101015). The samples were incubated at 37°C for 30-60 min with gentle inversion every 5-10 min. The digested cell suspension was filtered through a BD 40 μm cell strainer (Falcon, cat. no. 352340) 1-2 times. The filtrate was centrifuged at 300 × g for 5 min at 4°C. The cell pellet was resuspended in appropriate medium, mixed with an equal volume of red blood cell lysis buffer (Miltenyi, cat. no. 130-094-183), and incubated at 4°C for 10 min. After centrifugation at 300 × g for 5 min, the supernatant was removed. Finally, the cells were re-suspended with 1 ml RPMI 1640 medium (Conring, cat. no.10-040-CVR) with 0.04% BSA added. Single-cell suspension concentration and cell viability were then evaluated using the Luna-FL cell counter (Logos Biosystems, Korea) or the Trypan Blue staining method.

The freshly prepared single-cell suspension was adjusted to a concentration of 700–1200 cells/μL. Library preparation and loading were performed following the

---

[*]Tingjie Wang and Ruitao Long contributed equally to this work.

manufacturer's protocol for the MobiCube High-throughput Single Cell 3′ Transcriptome Set V2.1 (cat.no.PN-S050200301). The constructed libraries were sequenced on the DNBSEQ-T7 PE100 platform for high-throughput sequencing.

The FASTQ files were processed and aligned to the mouse GRCm39 genome using MobiVision software (V3.2) with unique molecular identifier (UMI) counts summarized for each barcode.

**Spatial transcriptomics sequencing**

Tumor tissues resected from ICC mouse tissues were immediately dissected, washed with 1 × PBS, and snap-frozen in liquid nitrogen-chilled isopentane. The specimens were embedded in OCT compound, placed on dry ice, and stored at -80°C. Sections of 10 μm thickness were mounted on chilled tissue optimization slides and gene expression slides (BMKMANU S1000). Each slide contains 1-8 identical 6.8×6.8 mm capture areas, comprising 2,000,000 spots with barcoded primers. These primers, attached at their 5' end, include a cleavage site, T7 promoter region, partial read 1 Illumina handle, unique spatial barcode, unique molecular identifier (UMI), and Poly(dT)VN sequence. The spots, with a diameter of 2.5 μm, are arranged in a centered regular hexagonal grid, and each spot is surrounded by six adjacent spots at a center-to-center distance of 4.8 μm. Tissue optimization, fixation, and H&E staining were conducted according to the manufacturer's protocol. Nuclei staining was also performed and imaged for cell segmentation. After reverse transcription and spatial library construction, libraries were quality-controlled and sequenced on the Illumina NovaSeq 6000 platform, producing 150 bp paired-end reads.

Four human ICC tissues were used for 10x HD spatial transcriptome (No. SHYJS-BC-2111001-06(GZ)). The Visium HD Spatial Gene Expression Platform (10x Genomics) achieves single-cell-level spatial resolution with 2 × 2 μm barcoded squares without gaps. The RNA quality of FFPE (formaldehyde-fixed, paraffin-embedded) tissue blocks is assessed by calculating the percentage of total RNA fragments >200nt (DV200) in the RNA extracted from tissue sections. For the high RNA quality, DV200 > 30% was required. According to the "Visium HD FFPE Tissue Preparation Handbook" (CG000684, 10x Genomics), a 5-μm-thick section of FFPE tissue was placed on a histology slide, dried at 42 °C for 3 h, dehydrated overnight, and incubated at 60 °C for 30 min. Subsequently, the tissue was deparaffinized following by hematoxylin-eosin (H&E) staining and imaging. Probe hybridization, probe ligation, slide preparation, probe release, extension, library construction were performed according to the "Visium HD Spatial Gene Expression Reagent Kits User Guide" (CG000685, 10x Genomics). The libraries were then sequenced on Illumina Novaseq 6000 with paired-end reads.

**Proteomics and phosphoproteomics sequencing**

Sample preparation for proteomics and phosphoproteomics. For PTM experiments, inhibitors were added to the lysis buffer (1% protease inhibitor cocktail, 1% phosphatase inhibitor for phosphorylation, 50 μM PR-619, 3 μM TSA and 50 mM NAM for acetylation). The remaining debris was removed by centrifugation at 12,000 g at 4 °C for 10 min. Finally, the supernatant was collected and the protein concentration was determined with BCA kit according to the manufacturer's instructions.

Trypsin digestion. The protein sample was added with 1 volume of pre-cooled acetone, vortexed to mix, and added with 4 volumes of pre-cooled acetone, precipitated at -20°C for 2 h. The precipitate was washed 2~3 times with the pre-cooled acetone. The protein sample was then redissolved in 200 mM TEAB and ultrasonically dispersed. Trypsin was added at 1:50 trypsin-to-protein mass ratio for the first digestion overnight. The sample was reduced with 5 mM dithiothreitol for 30 min at 56 °C and alkylated with 11 mM iodoacetamide for 15 min at room temperature in darkness. Finally, the peptides were desalted by Strata X SPE column.

Affinity Enrichment. Peptide mixtures were first incubated with IMAC microspheres suspension with vibration in loading buffer (50% acetonitrile/0.5% acetic acid). To remove the non-specifically adsorbed peptides, the IMAC microspheres were washed with 50% acetonitrile/0.5% acetic acid and 30% acetonitrile/0.1% trifluoroacetic acid, sequentially. To elute the enriched phosphopeptides, the elution buffer containing 10% NH OH was added and the enriched phosphopeptides were eluted with vibration. The supernatant containing phosphopeptides was collected and lyophilized for LC-MS/MS analysis.

LC-MS/MS Analysis. Mass Spectrometer: The tryptic peptides were dissolved in solvent A, directly loaded onto a home-made reversed-phase analytical column (25-cm length, 100 μm i.d.). The mobile phase consisted of solvent A (0.1% formic acid, 2% acetonitrile/in water) and solvent B (0.1% formic acid in acetonitrile). Peptides were separated with following gradient: 0-16 min, 2%-22%B; 16-22 min, 22%-35%B; 22-26 min, 35%-90%B; 26-30 min, 90%B, and all at a constant flow rate of 450 nl/minon a NanoElute UHPLC system (Bruker Daltonics). The peptides were subjected to capillary source separation followed by timsTOF Pro mass spectrometry. The electrospray voltage applied was 1.7 kV. Precursors and fragments were analyzed at the TOF detector. The timsTOF Pro was operated in data independent parallel accumulation serial fragmentation (dia-PASEF) mode. The full MS scan was set as100-1700（MS/MS scan range） and 22PASEF（MS/MS mode）-MS/MS scans were acquired per cycle. The MS/MS scan range was set as 395-1395 and isolation window was set as 20 m/z.

Database Search. The DIA data were processed using Spectronaut (v.18) software. Tandem mass spectra were searched against the Homo_sapiens_9606_SP_20231220.fasta (20429 entries) concatenated with reverse decoy database. Trypsin/P was specified as a cleavage enzyme, allowing up to 2 missing cleavages. Carbamidomethyl on Cys was specified as a fixed modification. Acetylation on protein N-terminal, oxidation on Met, and phosphorylation(S/T) were specified as variable modifications. False discovery rate (FDR) of protein, peptide and

PSM was adjusted to < 1%.

Bioinformatics analysis. Based on the quantitative results, fold changes (FC) between the two groups were calculated and p-values were computed using a t-test. Sites with $P < 0.05$ and log2FC > 1 were defined as differentially phosphorylated sites. Proteins harboring these differential sites were then subjected to GO and KEGG enrichment analysis.

**Single cell ATAC sequencing**

Nuclei preparation. The isolation, washing, and counting of nuclei suspensions were conducted in accordance with the established protocol: Nuclei Isolation for Single Cell ATAC Sequencing (10x Genomics; CG000169). The volume of chilled Diluted Nuclei Buffer (10x Genomics; 2000207) used for resuspension was determined based on the initial cell count and the desired final nuclei concentration. The resulting nuclei concentration was determined using a Countess II FL Automated Cell Counter. Subsequently, the nuclei were immediately utilized to generate single-cell ATAC-seq libraries.

Sequencing. The nuclei suspension was loaded into the Chromium Next GEM Chip H utilizing 10x reagents and barcoded with a Chromium Controller (10x Genomics). Subsequently, DNA fragments were amplified, and sequencing libraries were constructed utilizing reagents from the Chromium NextGEM Single Cell ATAC Reagent Kits v2 (10x Genomics), following the manufacturer's instructions. Furthermore, scATAC-seq libraries were prepared according to the Chromium NextGEM Single Cell ATAC Reagent Kits v2 User Guide Rev A (10x Genomics CG000496). The libraries were then pooled and sequenced on an Illumina NovaSeq platform with 2 × 150 paired-end configurations.

**Pathway enrichment analysis**

To explore the function of different cell types, we analyzed the gene enrichment in each cell type separately. Pathway enrichment was performed using the R package clusterProfiler (1) (3.14.3) in KEGG and GO database (Biological Process, BP). Pathways with a q-value of $< 0.05$ were selected for further analysis.

**Preparation of mouse bone marrow-derived macrophages (BMDMs)**

L929 cells were cultured to 100% confluence, after which the conditioned medium (L929-CM) was collected on days 3 and 5 and filtered through a 0.22 μm membrane. Bone marrow cells were isolated from 8-week-old mouse femurs and tibias under sterile conditions. After removing muscle tissue, bones were cut at both ends, and marrow was flushed out with medium. Cells were collected by centrifugation (500 × g, 10 min), treated with ice-cold RBC lysis buffer (1 min), and washed with two volumes of medium. After centrifugation, the pellet was resuspended in high-glucose DMEM supplemented with 10% FBS, 1% penicillin/streptomycin, and 30% L929-CM, then cultured in non-TC-treated dishes at 37°C with 5% $CO_2$. After 7 days, mature BMDMs were harvested using a cell scraper for subsequent experiments.

**Subcutaneous syngeneic allograft mouse model**

For tumor implantation, BR cells at 80% confluence were digested with trypsin, pelleted by centrifugation at 1,200 × g for 3 min, washed twice with PBS to remove serum residues, and resuspended in serum-free medium at $1 \times 10^7$ cells/mL. In subcutaneous tumor growth assays, $1 \times 10^6$ cells were injected into the flank of 8-week-old wild-type FVB mice. For BMDM/BR co-injection experiments, FVB mice received subcutaneous injections of either $1 \times 10^6$ BR cells alone or a mixture of $1 \times 10^6$ BR cells and $1 \times 10^6$ BMDMs into contralateral sites.

Tumor growth was monitored every two days by measuring the longest and shortest diameters, and tumor volume was calculated as (length × width$^2$) / 2. All animal procedures were conducted in accordance with institutional animal care guidelines, and mice were euthanized before the tumor volume reached 1,500 mm$^3$ for subsequent analyses.

**Isolation of tumor-infiltrating leukocytes**

Tumor tissues were minced into approximately 1 mm$^3$ pieces and digested in high-glucose DMEM containing collagenase IV (1 mg/mL), hyaluronidase (0.25 mg/mL), and DNase I (0.1 mg/mL). The digestion was performed at 37°C with shaking at 150 rpm for 3 h. The digested mixture was passed through a 70 μm cell strainer to obtain a single-cell suspension. Cells were collected by centrifugation at 500 × g for 5 min, then resuspended in 40% Percoll and layered over 80% Percoll to form a two-layer gradient. The gradient was centrifuged at 800 × g for 30 min at room temperature without a brake. The interface layer containing tumor-infiltrating leukocytes was carefully collected for subsequent analyses.

**Combined Thbs1/OSM blockade in ICC Mouse Models**

A combination of genetic knockout of Thbs1 and pharmacological inhibition of OSM was conducted in vivo. A primary ICC model was established in Thbs1$^{fl/fl}$ and Thbs1$^{fl/fl}$;Lyz2-Cre mice via hydrodynamic tail vein injection of AKT1 and NICD1 overexpression plasmids (designated as Day 0). Beginning on day 10 post-modeling, mice were randomly assigned to one of four treatment groups: (1) Thbs1$^{fl/fl}$ + vehicle control; (2) Thbs1$^{fl/fl}$ + OSM-SMI-10B; (3) Thbs1$^{fl/fl}$;Lyz2-Cre + vehicle control; (4) Thbs1$^{fl/fl}$;Lyz2-Cre + OSM-SMI-10B.

The OSM-SMI-10B compound was dissolved in a 20% (w/v) sulfobutylether-β-cyclodextrin (SBE-β-CD) saline solution to achieve a working concentration of 20 μg/mL. All drug-treated groups received intraperitoneal injections of OSM-SMI-10B at a dose of 20 mg/kg every two days. The vehicle control groups were administered an equivalent volume of the 20% SBE-β-CD solution on the same schedule. The experiment was terminated on day 31 post-modeling. Liver phenotypes were documented, and tissues were collected for subsequent analysis.

**Flow cytometry**

For each tumor sample, approximately $1 \times 10^6$ tumor-infiltrating leukocytes were collected by centrifugation and resuspended in 100 μL PBS for staining. Cells were stained with mouse CD45-APC, mouse F4/80-PerCP-Cy5.5, and mouse/human cross-reactive CD11b-FITC antibodies at 4°C for 30 min in the dark. After staining, cells were washed with PBS, filtered through a 200-mesh cell strainer, and analyzed using a BD Accuri™ C6 flow cytometer.

For BMDMs, surface staining was performed using F4/80 and CD11b antibodies. Cells were then fixed with Fixation Buffer at room temperature in the dark for 20 min. After fixation, cells were permeabilized using Intracellular Staining Permeabilization Wash Buffer (1×) and stained with mouse CD206-APC antibody at 4°C for 30 min in the dark. Following washing and resuspension, samples were analyzed by flow cytometry on the BD Accuri™ C6. Antibody information for flow cytometry is detailed in Supplementary Table S6.

**Western blotting**

Cells were lysed on ice by sonication in RIPA buffer containing protease and phosphatase inhibitors, or tissues were homogenized on ice in the same buffer. Lysates were centrifuged at 12,000 rpm for 10 min at 4°C to collect the supernatant. Protein concentration was determined using a BCA assay. Samples were mixed with reducing loading buffer and denatured at 95°C for 10 min in a metal bath. Equal amounts of protein (30 μg) and prestained protein markers were loaded onto SDS-PAGE gels. Electrophoresis was performed at 30 V constant voltage for 20 min, then at 120 V for approximately 2 h until sufficient separation was achieved. PVDF membranes were activated with ethanol and assembled into transfer sandwiches. Proteins were transferred in semi-dry transfer buffer (NCM biotech) at a constant current of 400 mA for 30 min. Membranes were blocked at room temperature on a shaker for 1 h with 5% BSA in TBST for phosphorylated proteins or 5% non-fat milk in TBST for other proteins. Primary antibodies were incubated overnight at 4°C. After washing three times with TBST for 5 min each, membranes were incubated with HRP-conjugated secondary antibodies for 1 h at room temperature. After final washes, enhanced chemiluminescence (ECL) substrate was applied, and signals were detected using the GeneSys imaging system. Appropriate loading controls (GAPDH, β-Actin, or α-Tubulin) were selected based on confirmed stable expression under the specific experimental treatments. The specific control used for each experiment is clearly indicated in the figures. Antibody information for Western blotting is detailed in Supplementary Table S6.

**Immunohistochemistry (IHC)**

Paraffin-embedded tissue sections were deparaffinized in xylene and graded ethanol series. Antigen retrieval was performed in sodium citrate buffer using microwave heating (medium power 11 min, rest 8 min, low power 9 min), then cooled to room temperature. For nuclear antigens, the medium power heating was extended to 25 min. Endogenous peroxidase was quenched with 3% hydrogen peroxide at room temperature for 25 min in the dark. Sections were blocked with goat serum for 30 min at room temperature, then incubated with primary antibodies overnight at 4°C in a humidified chamber. After washing, sections were incubated with secondary antibodies diluted 1:1000 in TBST at room temperature for 2 h, followed by DAB development. Antibody information for IHC is detailed in Supplementary Table S6.

**Multiplex immunofluorescence (mIF)**

Sections were pretreated as in IHC. Sequential incubations of primary and HRP-conjugated secondary antibodies were performed, followed by TSA fluorophore incubation at 37°C for 10 min in the dark. Between each staining cycle, antigen retrieval was conducted at 95°C for 15 min. This cycle was repeated until all targets were stained. Nuclei were counterstained with DAPI, and sections were mounted with antifade reagent before fluorescence imaging. Antibody information for mIF is detailed in Supplementary Table S6.

**Quantitative RT-PCR (qRT-PCR)**

Total RNA was extracted using TRIzol reagent and quantified by NanoDrop. cDNA was synthesized from 2 μg RNA using Goldenstar™ RT6 Kit (Tsingke) with gDNA removal (42°C 2 min, 60°C 5 min) followed by reverse transcription (50°C 15 min, 85°C 5 min). cDNA was diluted 10-fold before use. Quantitative PCR was performed in 20 μL reactions containing 4 μL diluted cDNA, 0.4 μM of each forward and reverse primer, and SYBR Green Master Mix （Tsingke） on a LightCycler® 96 system (Roche) with cycling conditions: 95°C for 10 min; 40 cycles of 95°C for 15 s, 60°C for 15 s, and 72°C for 15 s. Relative expression was calculated by the 2−ΔΔCt method using β-actin for normalization. The sequences of all primers used for qRT-PCR are detailed in Supplementary Table S7.

**siRNA interference**

siRNA transfection was performed using Lipofectamine 3000 ((Thermo Fisher Scientific). Tumor cells at 50% confluence in 6-well plates were transfected with Complex 1 (125 μL Opti-MEM + 5 μL Lipofectamine 3000) and Complex 2 (125 μL Opti-MEM + 10 μL 20 μM siRNA), which were mixed and incubated at room temperature for 10 – 15 min before being added to cells in 1.75 mL complete medium. BMDMs (1 × $10^6$ cells/well) were seeded in untreated 12-well plates and transfected with Complex 1 (75 μL Opti-MEM + 75 μL Lipofectamine 3000) and Complex 2 (60 μL Opti-MEM + 7 μL 20 μM siRNA), mixed and incubated for 10 – 15 min at room temperature, then added to cells cultured in 850 μL HG-DMEM with 5% FBS.Cells were harvested 24 h post-transfection for co-culture experiments and 48 h post-transfection for qPCR and Western blot analyses. The sequences of all siRNAs used in this study are detailed in Supplementary Table S8.

**Lentiviral transduction and stable cell line generation**

BR cells were seeded in 12-well plates and allowed to grow to 20% confluence. The LV3-shNC/LV3-shCD47 lentiviruses (GenePharma) were diluted 1:10 in complete RPMI-1640 medium containing 5 μg/mL Polybrene for 24-hour transduction. After infection, the medium was replaced with fresh complete medium, followed by selection with 2 μg/mL puromycin until complete death of wild-type cells was observed. The surviving cells were collected for CD47 knockdown efficiency validation by Western blot and qRT-PCR analyses. The sequences of shRNA used in this study are detailed in Supplementary Table S9.

**Transwell migration assay**

A 24-well Transwell system with 8 μm pore size was used. Tumor cells suspended in serum-free medium were seeded in the upper chamber. Depending on the experimental design, the lower chamber was either seeded with BMDMs or left cell-free and was filled with complete medium alone or supplemented with recombinant proteins, inhibitors, or a combination of both. After incubation for 24 h in an 37°C incubator, the upper chamber was carefully removed, rinsed with PBS, and fixed with 4% paraformaldehyde for 15 min. Cells on the upper surface were gently wiped off with a PBS-moistened cotton swab. Migrated cells on the lower surface were stained with 0.1% crystal violet for 15 min, washed with distilled water, and imaged under a microscope.

**Scratch wound migration assay**

Log-phase ICC cells were seeded in 6-well plates until reaching 90% confluence. Three parallel scratches were made with a 10 μL pipette tip, followed by PBS washing. Experimental groups were treated with 1% FBS medium containing 10 ng/mL Thbs1, while control groups received 1% FBS medium only. Wound closure was monitored and photographed using bright-field microscopy at 0, 6, 12, 24, 36, 48, and 72 h for quantitative analysis.

**Contact co-culture**

Tumor cells and macrophages were prepared as single-cell suspensions and counted. Cells were mixed at a 1:1 ratio and seeded into 6-well plates. Macrophages cultured alone served as the control. For co-culture, macrophages and tumor cells were incubated together at 37°C with 5% $CO_2$ for 48 h, after which all cells were collected for qRT-PCR analysis of polarization markers.

**Transwell co-culture assay**

A 12-well Transwell system with 0.4 μm pores was used. Tumor cells were seeded in the lower chamber and allowed to adhere for 12 h before carefully placing the upper chamber containing macrophages. After 24 or 48 h of co-culture, cells from both chambers were collected for qRT-PCR and Western blot analyses. Additionally, EdU incorporation assays were performed on lower chamber cells, and flow cytometry was conducted on upper chamber cells.

**Macrophage-induced tumor cell migration assay**

A 24-well Transwell system with 8 μm pores was used. Macrophages were seeded in the lower chamber and allowed to adhere. Tumor cells suspended in serum-free medium were carefully seeded in the upper chamber. Co-culture was carried out at 37°C for 24 h, followed by staining and imaging of the upper chamber according to the cell migration assay protocol.

**Tumor conditioned medium (TCM)-induced macrophage polarization**

ICC cells were cultured to approximately 90% confluence and then cultured for an additional 48 h to collect TCM filtered through a 0.45 μm membrane. For polarization induction, TCM and complete HG-DMEM medium were mixed at a 3:1 ratio and added to macrophages; controls received complete HG-DMEM only. Cells were collected at various time points post-treatment for qRT-PCR or flow cytometry analysis.

**Colony formation assay**

Cells (500/well) were seeded in 6-well plates and cultured for 2 weeks with medium change every 4 days. When colonies became visible, cells were washed with PBS, fixed with 4% paraformaldehyde for 15 min, stained with 0.1% crystal violet for 30 min at room temperature, and rinsed with distilled water. Colony images were captured for quantitative analysis.

**EdU incorporation assay**

Cells were incubated with 10 μM EdU (Beyotime) in complete medium (1 h for BR; 2 h for HuCCT1, RBE andQBC939) in an incubator. After fixation (4% PFA, 15 min) and permeabilization (0.3% Triton X-100, 15 min), the incorporated EdU was then fluorescently labeled via Click reaction after 30 min incubation in dark (Beyotime Kit C0075,), followed by nuclear counterstaining with DAPI (5 μg/mL, 10 min). Fluorescence images were acquired using an inverted microscope (555 nm/405 nm) and analyzed with ImageJ. EdU positive rate (%) = (EdU positive cells / DAPI positive cells) × 100% (5 fields pergroup).

**CCK-8 assay**

Cells ($2 \times 10^3$/well) were seeded in 96-well plates with 100 μL medium. After treatment, medium was replaced with 90 μL fresh medium containing 10 μL CCK-8 reagent (Abbkine), incubated at 37°C for 1 h protected from light, then absorbance was measured at 450 nm using a microplate reader.

**mIHC image acquisition and quantification**

During image acquisition, an eight-channel fluorescence digital slide scanner (AF-KL-20-8, AiFang Biological, China) was used to capture images of mIHC slides, which were saved in the kfbf format. In the image data analysis stage, custom algorithms were developed in the VISIOPHARM software (VISIOPHARM, Hoersholm, Denmark) to analyze the images. To ensure consistent analysis results, fixed, uniform threshold parameters were set in the VISIOPHARM application to identify and quantify cell positivity rates and spatial infiltration. After the analysis was completed, the results were exported. After data collection, GraphPad Prism 8.0.2 was used to analyze the data. We calculated the densities of THBS1-, and CD68-positive cells as the number of positive cells per $mm^2$ within the tumor tissues. Then, we performed survival analyses in R using the survival package (v3.5-7). Patients were dichotomized into high and low groups using the cohort mean as the cutoff, Kaplan–Meier curves were generated, and differences between groups were assessed with the log-rank test to obtain P values.

**Quantification and statistical analysis**

All experiments were conducted with rigorous reproducibility, including at least three independent biological replicates per group ($n \geq 3$) to ensure result stability and reliability. Technical replicates were performed for each sample to minimize experimental error. For animal studies, each group included no fewer than three animals ($n \geq 3$), all treated and sampled under identical conditions.

Statistical analyses were performed using GraphPad Prism or R software. Comparisons between two groups were conducted using two-tailed unpaired Student's t-tests. For multiple group comparisons, Statistical significance was analyzed by one-way ANOVA ($p < 0.05$) followed by Dunnett's multiple comparisons test for pairwise group comparisons. Significance levels are indicated as follows: *$p < 0.05$, **$p < 0.01$, ***$p < 0.001$, ****$p < 0.0001$, and ns: not significant. All data are representative of three biologically independent experiments, unless otherwise indicated.

### Supplementary Figures

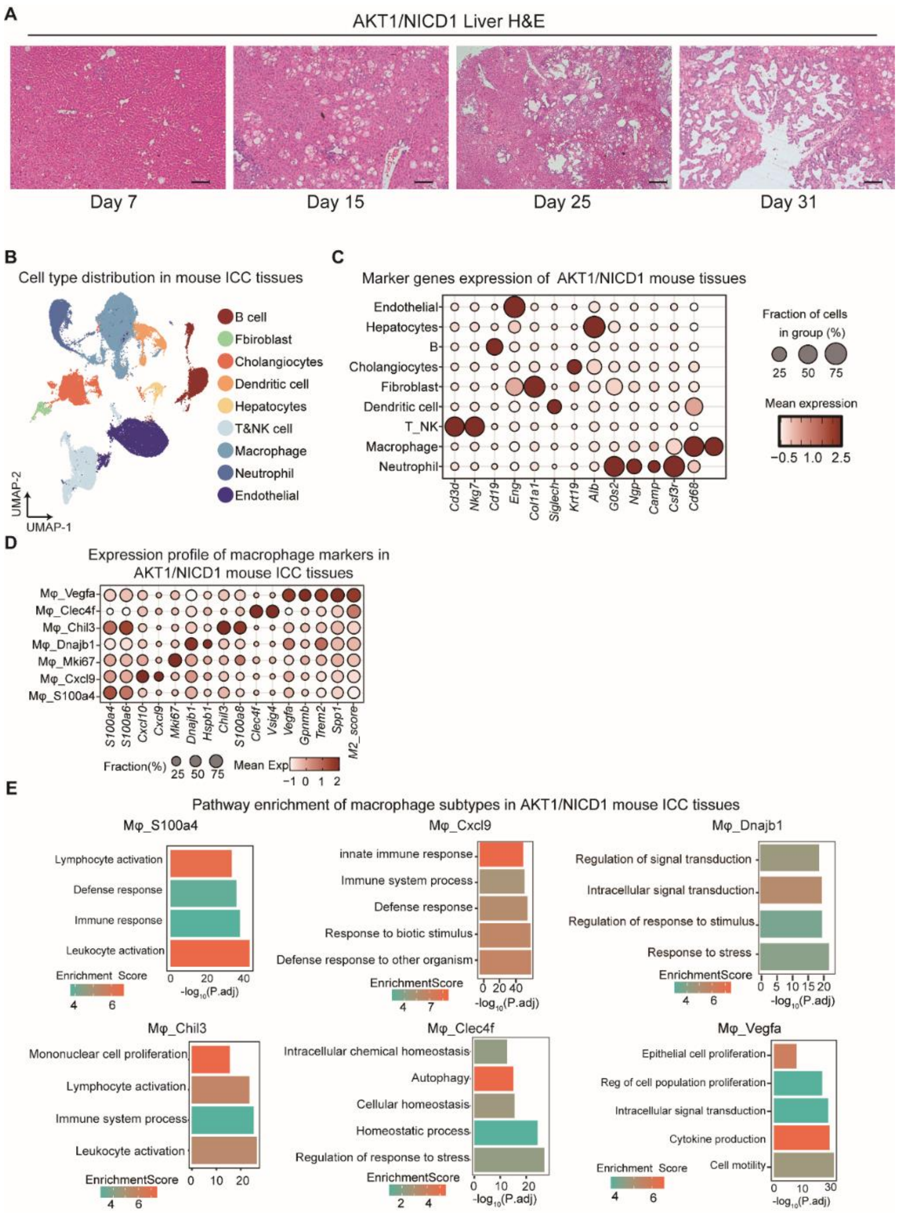

**Figure S1. Cell profiles and functional enrichment in the AKT1/NICD1 mouse ICC tissues. (A)** Representative H&E-stained liver sections at days 7, 15, 25, and 31 post-hydrodynamic tail vein injection of AKT1/NICD1. Scale bar, 50 μm. **(B)** Cell type distribution in AKT1/NICD1 mouse tissues. UMAP plot visualizes the cell types. Bubble plot showing the expression profiles of representative marker genes of major cell types **(C)** and macrophage subtypes **(D)** in the AKT1/NICD1 mouse ICC tissues. **(E)** Bar plot showing the pathway enrichment of the macrophage subclusters in the AKT1/NICD1 mouse ICC tissues. The X-axis represents enrichment scores.

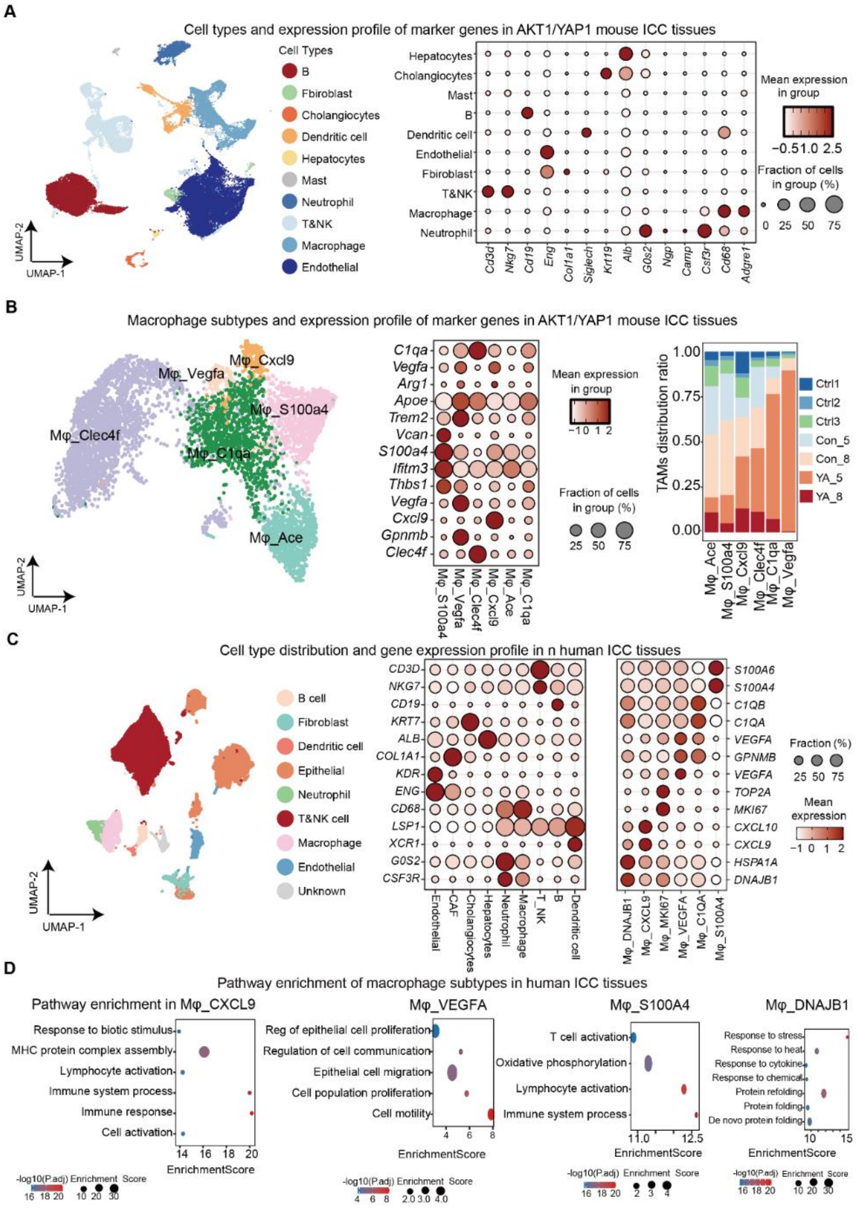

**Figure S2. Cell profiles and functional enrichment in the AKT1/YAP1 mouse and human ICC tissues. (A)** UMAP plot visualizes cell type distribution in AKT1/YAP1 mouse tissues, and bubble plot showing the expression profiles of representative marker genes of major cell types. **(B)** UMAP plot visualizes macrophage subtypes (left), and bubble plot showing the expression profiles of representative marker genes of macrophage subtypes in the AKT1/YAP1 mouse ICC tissues, and bar plot showing the macrophage subtypes distribution of the tumor progression in the AKT1/YAP1 mouse ICC tissues (right). Ctrl, empty vectors with HSB2. Con_5, harvested 5 weeks after empty vectors with HSB2. Con_8, harvested 8 weeks after empty vectors with HSB2. YA_5, harvested 5 weeks after AKT1/YAP1 injection. YA_8, harvested 8 weeks after AKT1/YAP1 injection. **(C)** UMAP and bubble plot showing the cell types and marker genes profile in the human ICC tissues. (**D)** Dot plot showing the enriched pathways in the human macrophage subtypes.

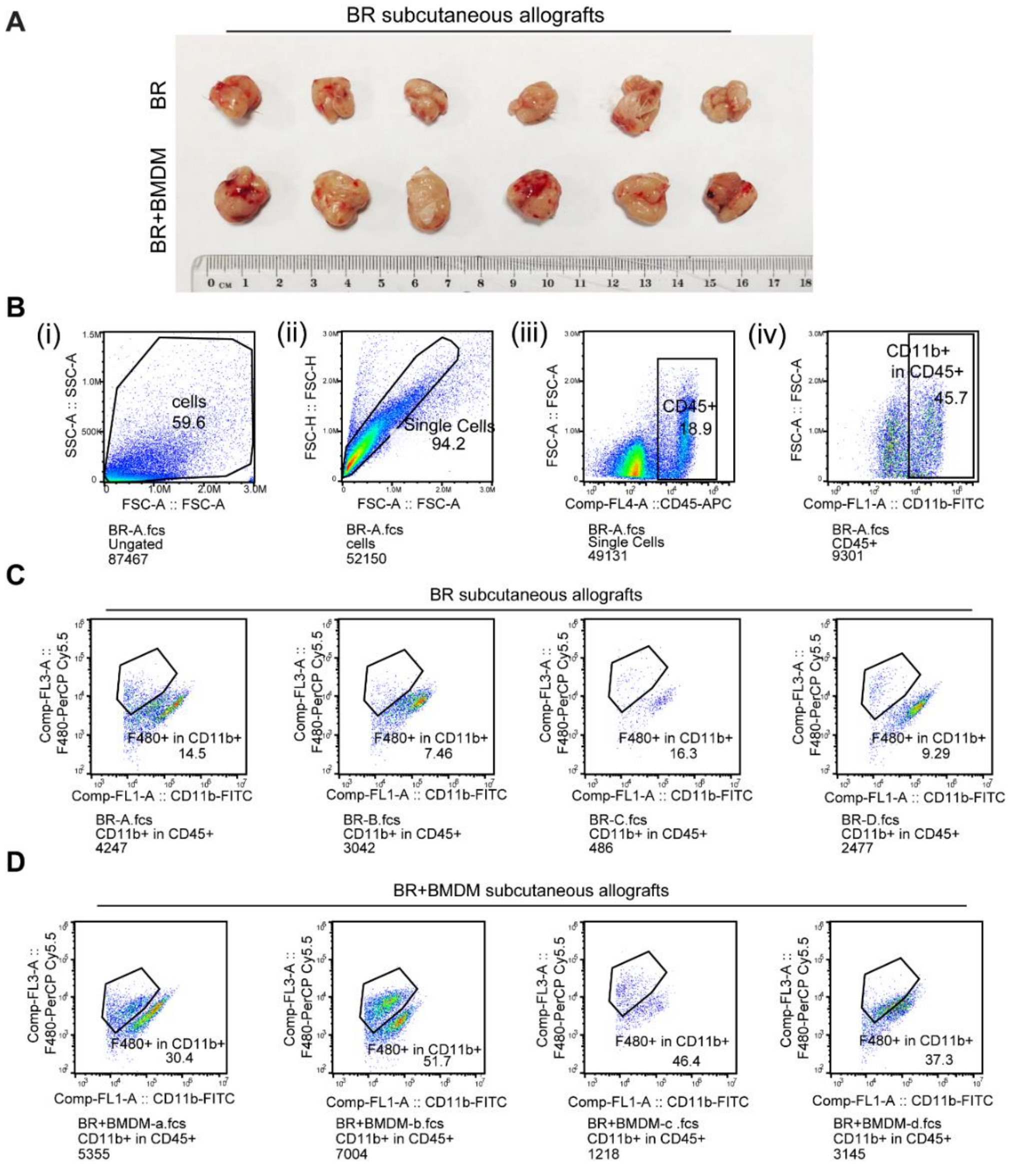

**Figure S3. Co-injection of BMDMs with BR cells enhanced macrophage infiltration and promoted the growth of ICC allografts. (A)** Images of subcutaneous allografts from the BR cell alone and the BR/BMDM co-injection groups. n = 6 mice. **(B)** Quantification of tumor-infiltrating macrophages (F4/80$^+$CD11b$^+$ cells) by flow cytometry. Data are presented as mean ± SD; Significance was determined by a two-tailed unpaired t-test. $**p < 0.01$, and n = 4 mice. **(C)** Flow cytometry gating strategy for macrophage co-injection allograft experiments. Cells were first gated based on forward scatter area (FSC-A) and side scatter area (SSC-A) to select the cell population, followed by gating on single cells using FSC-A versus FSC-H. CD45$^+$ cells were then gated using FL4-A versus FSC-A, and subsequently CD11b$^+$ cells were selected within the CD45$^+$ population. The proportion of F4/80$^+$ cells was finally analyzed within the CD11b$^+$ subset. **(D)** Percentage of F4/80$^+$ cells within the CD11b$^+$ population in ICC allografts derived from mice injected with BR cells alone (control group).

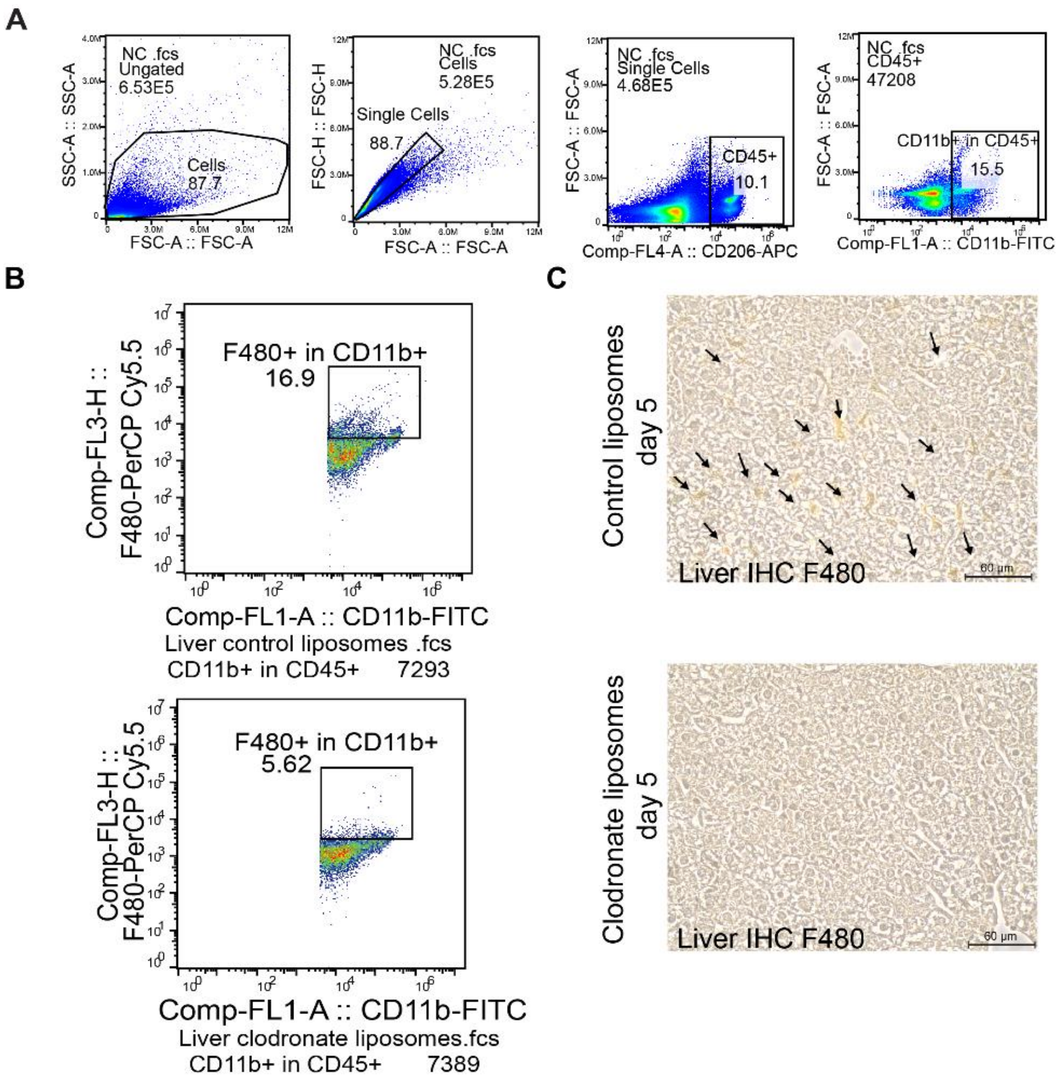

**Figure S4. Clodronate liposome-mediated depletion of liver macrophages effectively inhibited the progression of AKT1/NICD1-induced ICC. (A)** Flow cytometry gating strategy for macrophage depletion experiments. Cells were gated sequentially as follows: FSC-A versus SSC-A for cell population (the first), FSC-A versus FSC-H for single cells (the second), FL4-A versus FSC-A for $CD45^+$ cells (the third), and $CD11b^+$ cells within the $CD45^+$ population (the fourth). **(B)** Flow cytometry analysis of liver tissues from wild-type mice 5 days after injection of clodronate liposomes or control liposomes. The percentage of macrophage infiltration ($F4/80^+CD11b^+$ cells) among total live cells is shown. **(C)** Representative images of F4/80 IHC staining on paraffin-embedded liver sections from wild-type mice 5 days after injection of clodronate liposomes or control liposomes. Arrows indicate F4/80-positive cells. Scale bar = 60 μm.

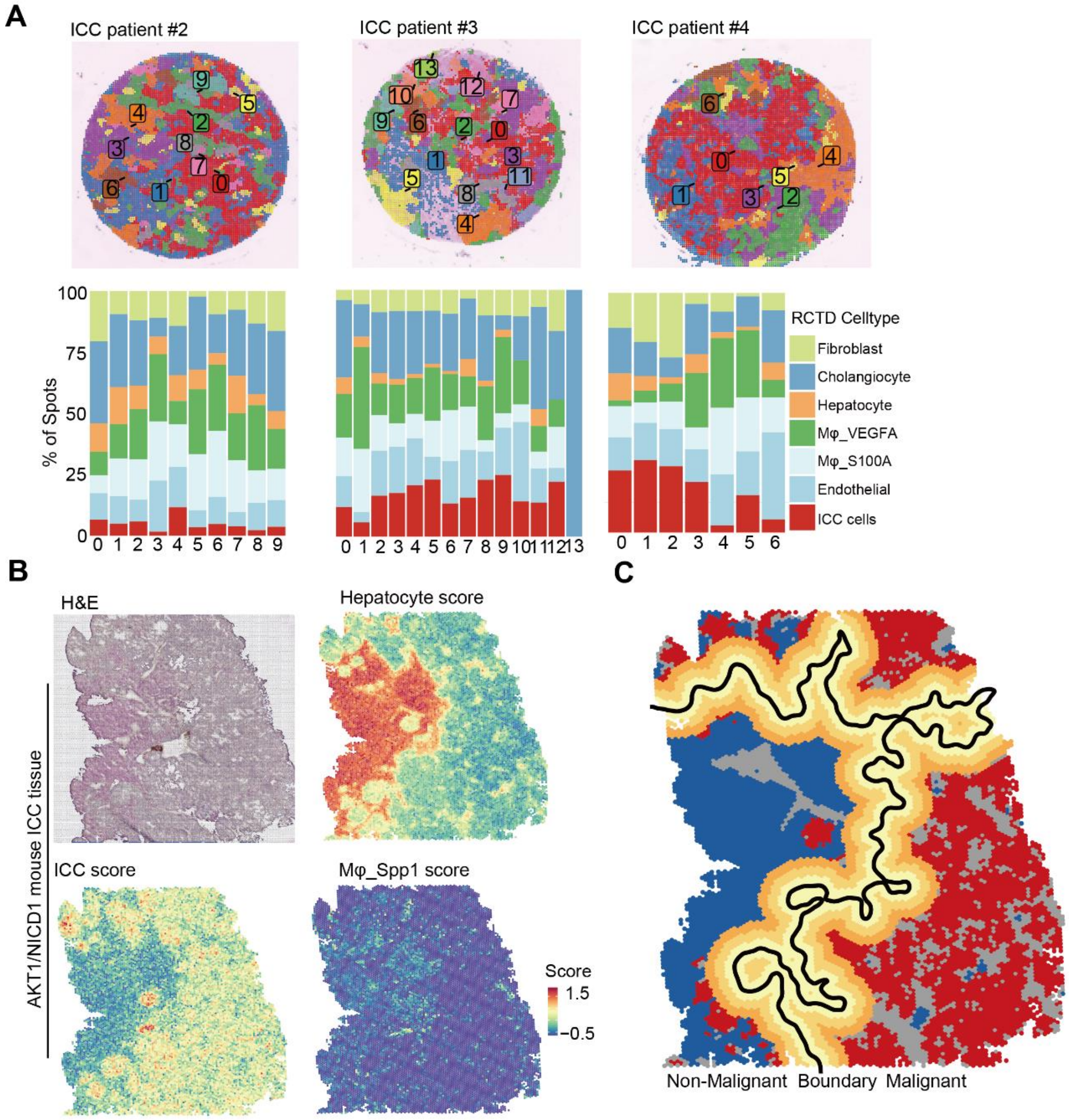

**Figure S5. Spatial distribution of Mφ_VEGFA and its spatial proximity to ICC cells. (A)** BANKSY spatial domains and their cellular celltype composition via RCTD analysis in the three ICC patient tissues (Patient 2#, 3#, and 4#). **(B)** H&E and heatmap showing the distribution of cell types in the AKT1/NICD1 ICC tissues. **(C)** Spatial segmentation of AKT1/NICD1 mouse ICC tissue.

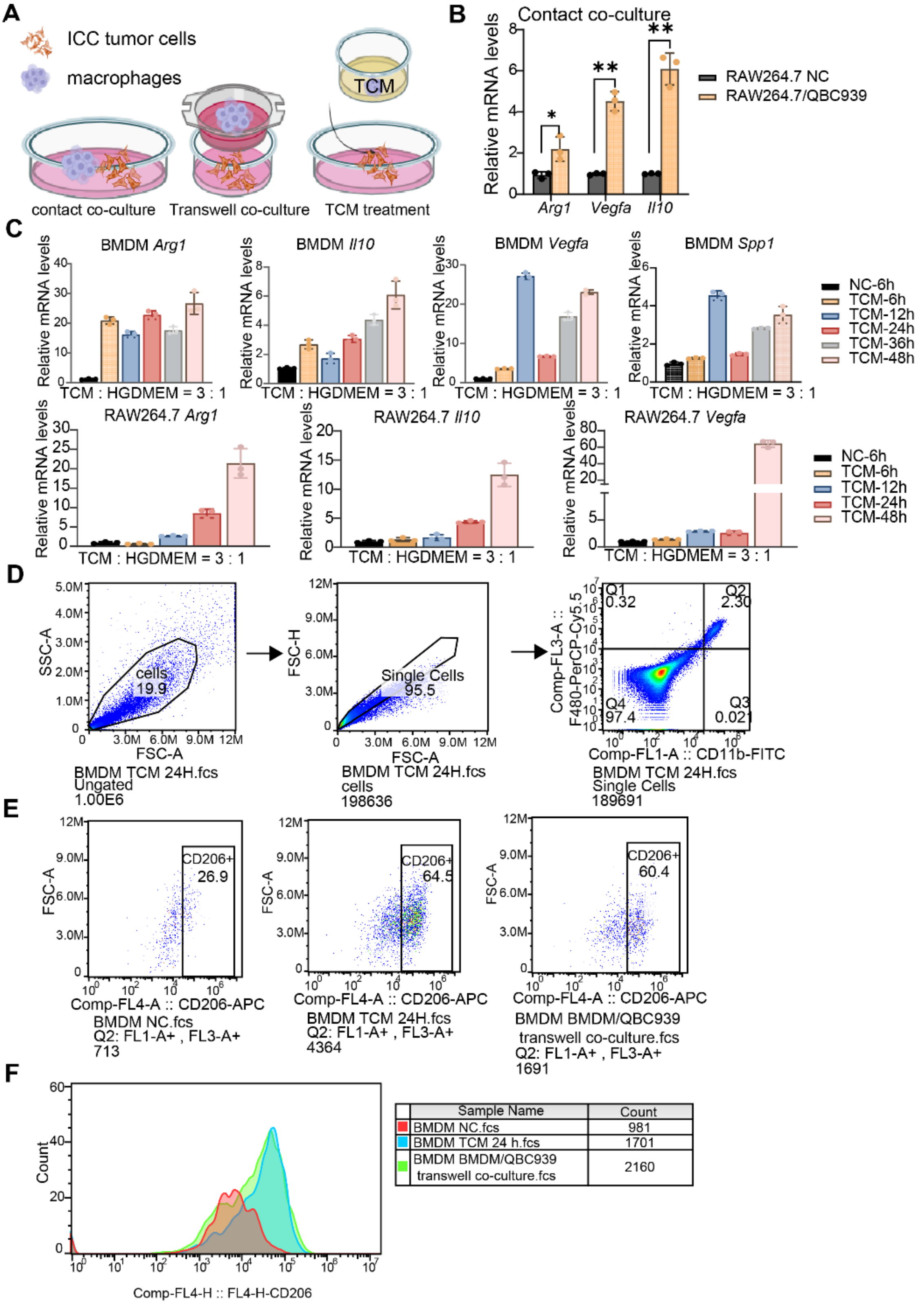

| Sample Name | Count |
|---|---|
| BMDM NC.fcs | 981 |
| BMDM TCM 24 h.fcs | 1701 |
| BMDM BMDM/QBC939 transwell co-culture.fcs | 2160 |

**Figure S6. ICC/Macrophage co-culture drives M2-like polarization assessed by qRT-PCR and flow cytometry. (A)** Schematic diagram of three macrophage polarization models: direct contact co-culture, transwell co-culture, and culture with tumor-conditioned medium (TCM). **(B)** qRT-PCR analysis of *Arg1*, *IL-10*, and *Vegfa* mRNA levels in a direct contact co-culture system of QBC939 and RAW264.7 cells at a 1:1 ratio for 36 h, with mixed harvesting of both cell types for RNA extraction. Data are presented as mean ± SD. Significance was determined by a two-tailed unpaired t-test (*$p$ < 0.05, and **$p$ < 0.01). **(C)** BMDMs (top) or RAW264.7 cells (bottom) were treated with fresh medium diluted 1:3 with TCM collected from ICC cell lines; cells were harvested at indicated time points for qRT-PCR analysis of *Arg1*, *IL10*, and *Vegfa* mRNA levels. **(D)** Flow cytometry gating strategy for BMDM polarization assays. Cells were gated based on FSC-A versus SSC-A to select the viable cell population, followed by FSC-A versus FSC-H to select single cells. Mature macrophages were identified by F4/80$^+$CD11b$^+$ double-positive gating. **(E)** Flow cytometry analysis of CD206 expression in BMDMs (the first), BMDMs TCM-treated for 24 h (the second), and BMDMs transwell co-cultured with QBC939 for 48 h (the third). **(F)** Histogram showing fluorescence intensity distribution of CD206 expression in BMDMs (red), BMDMs TCM-treated for 24 h (blue), and BMDMs transwell co-cultured with QBC939 for 48 h (green).

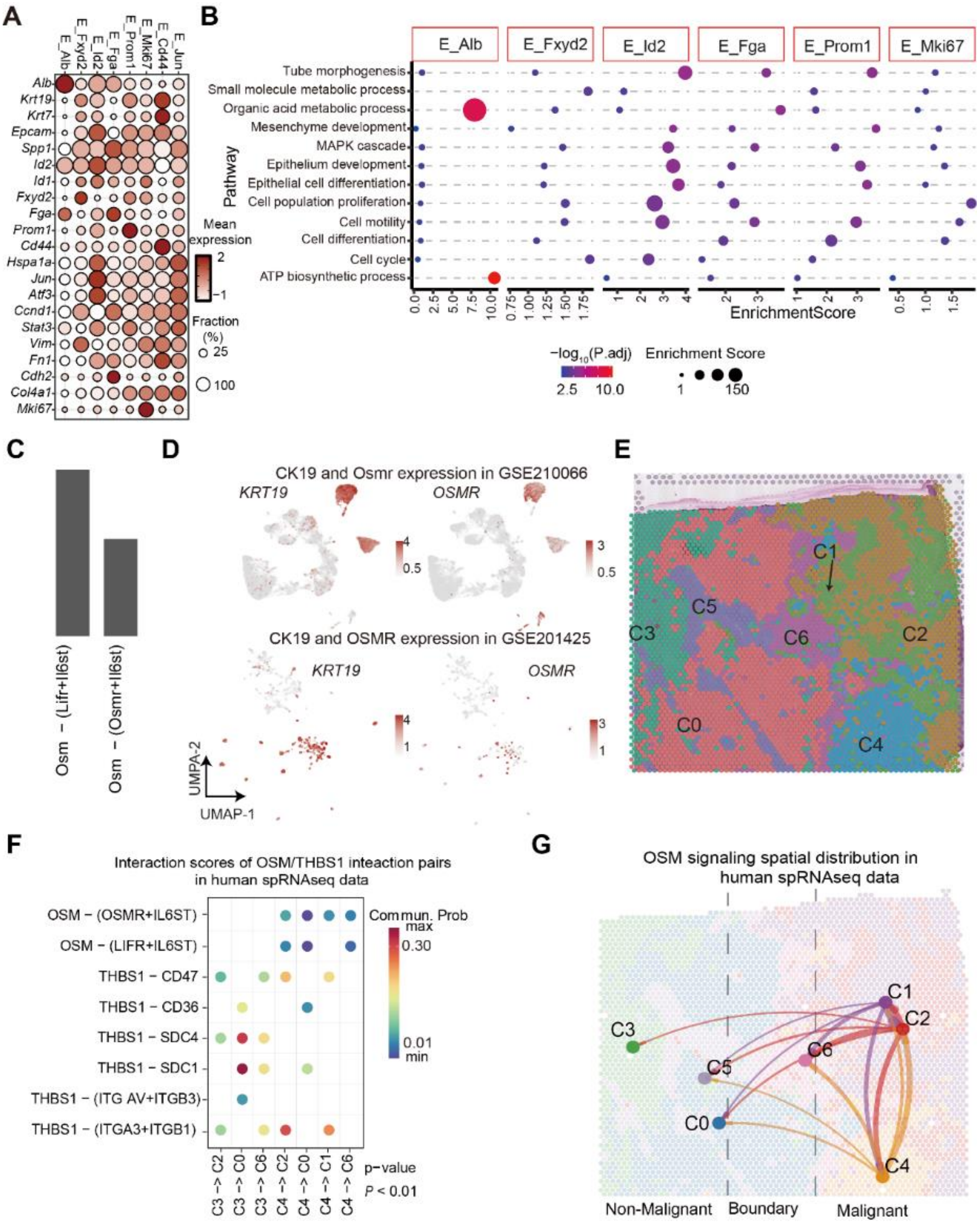

**Figure S7 OSM-OSMR cellular interaction in AKT1/NICD1 mouse and human ICC tissues. (A)** Bubble plot showing the expression patterns of representative marker genes for epithelial subclusters in human ICC tissues. The X-axis indicates marker genes, and the Y-axis denotes epithelial subclusters. The color gradient from white to red represents increasing expression levels, while bubble size reflects the proportion of cells expressing each gene. **(B)** Dot plot showing the pathway enrichment results in the epithelial subtypes. **(C)** Interaction strength of ligand and its receptor in the AKT1/NICD1 mouse ICC tissues. **(D)** Expression of OSMR in the human ICC tissues. **(E)** Spatial clustering results of human ICC tissue. **(F)** Bubble plot maps spatially resolved cell-cell communication networks mediated by *OSM* ligands in spatial transcriptomic data. Rows denote interacting cell clusters, while columns represent ligand-receptor interaction pairs. Bubble size and color intensity reflect interaction strength. **(G)** Spatial distribution patterns of the OSM signaling axis across tissue compartments: non-malignant adjacent, boundary, and malignant tumor regions.

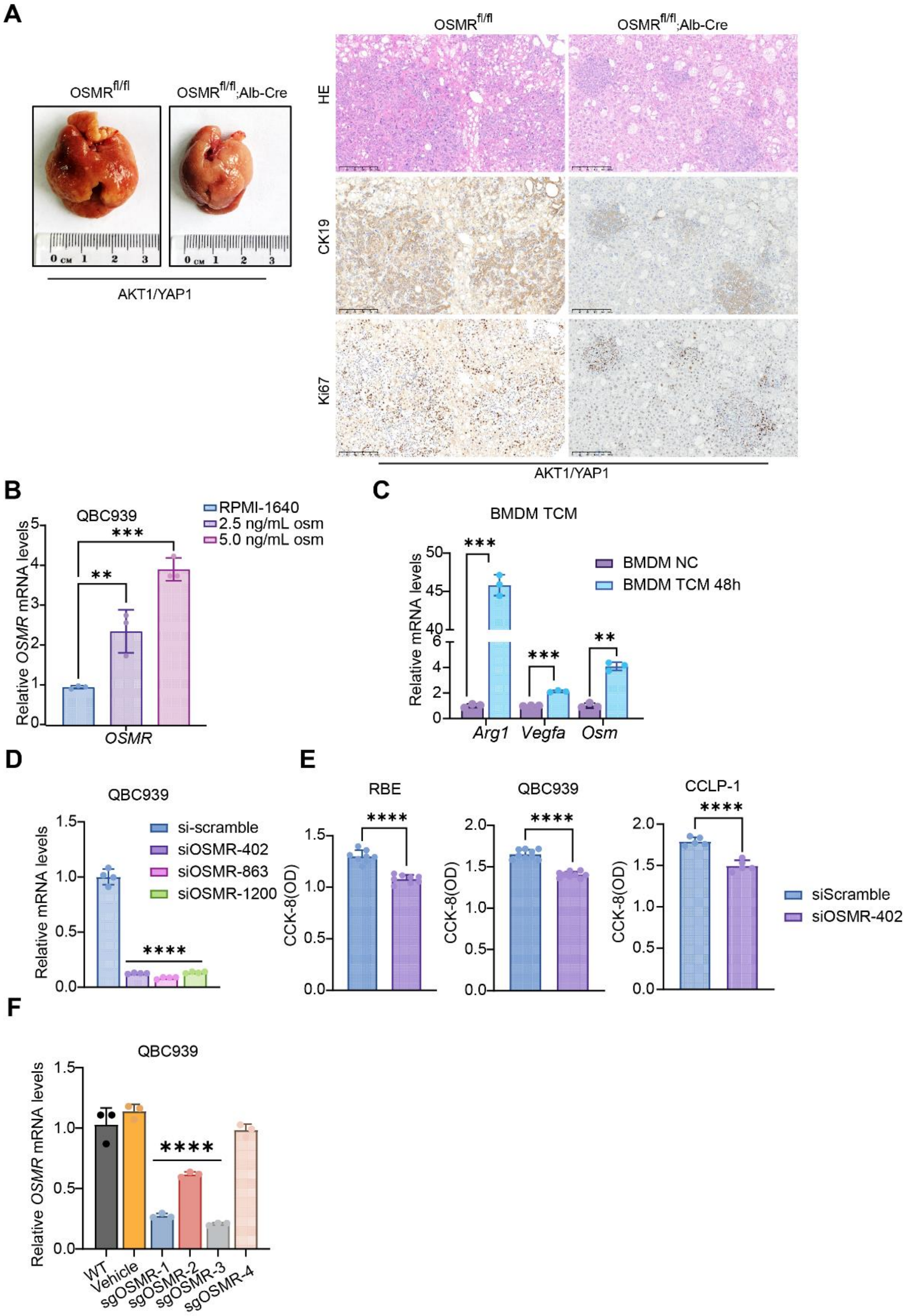

**Figure S8 Macrophage-derived OSM upregulates OSMR to promote ICC cell growth.** **(A)** Representative liver images from $Osmr^{fl/fl}$ and $Osmr^{fl/fl}$;Alb-Cre mice injected with AKT1/YAP1**;** Immunohistochemical staining of CK19, Ki67, and HE staining in paraffin-embedded liver sections from $Osmr^{fl/fl}$ and $Osmr^{fl/fl}$;Alb-Cre mice injected with AKT1/YAP1. Scale bars, 200 μm. **(B)** qRT-PCR analysis of *OSMR* transcript levels in QBC939 cells treated with RPMI-1640 complete medium (control), 2.5 ng/mL, or 5.0 ng/mL recombinant OSM protein. **(C)** Quantitative RT-PCR analysis of *Arg1*, *Vegfa*, and *Osm* transcript levels in BMDMs treated with TCM for 24 h. M0 served as control. **(D)** qRT-PCR analysis of OSMR transcript levels in QBC939 cells transfected with siScramble or siOSMR sequences (402, 863, and 1200). **(E)** CCK-8 assay measuring optical density in ICC cell lines QBC939, RBE, and CCLP-1 transfected with siScramble or siOSMR402 after 48 h. **(F)** qRT-PCR analysis of *OSMR* transcript levels in QBC939 cells infected with none, lenti-virus with blank vector (vehicle), or lenti-virus with sgOSMR sequences (sgOSMR-1, -2, -3, -4). Data are presented as mean ± SD. Statistical significance was analyzed by one-way ANOVA ($p$ < 0.05) followed by Dunnett's test (p < 0.05) followed by Dunnett's test(****$p$ < 0.0001).

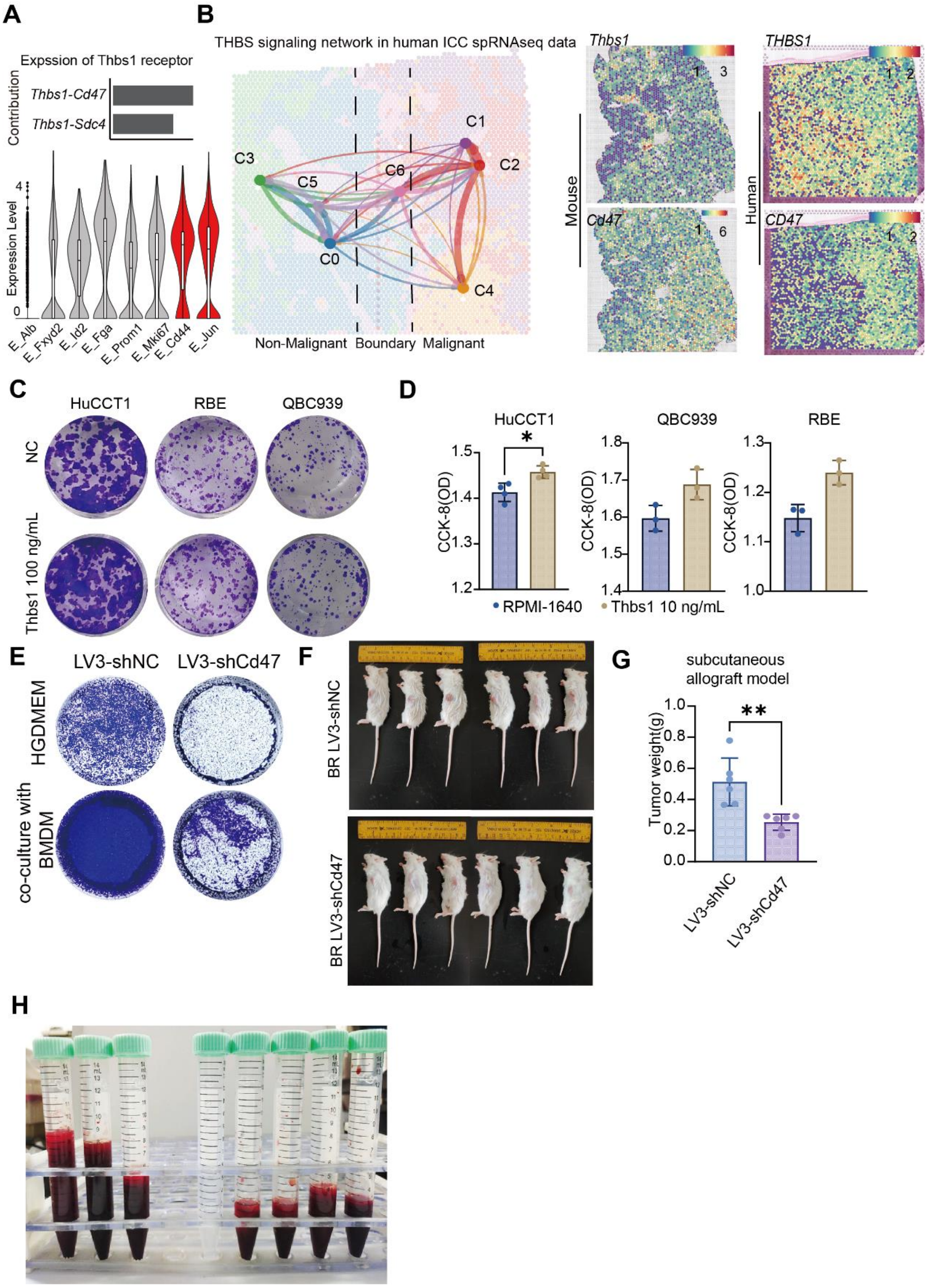

**Figure S9. THBS1 downstream signaling in human ICC and AKT1/NICD1 mouse ICC subtypes. (A)** Interaction strength of ligand Thbs1 and its receptors (top) and their expression level (bottom) in the AKT1/NICD1 mouse ICC tissues. **(B)** The interaction intensity of the THBS1 signaling axis between subpopulations in the human ICC spatial transcriptomics data, where C0 and C1 represent ICC cell-dominant subpopulations, and C5 and C6 represent macrophage-dominant subpopulations, with C5 and C6 localized at the ICC cell periphery (left). Spatial expression patterns of the *THBS1-CD47* interacting gene pair in the AKT1/NICD1 mouse ICC and human ICC tissues. The color gradient from blue to red indicates increasing expression levels (right). **(C)** Representative crystal violet–stained images for colony formation assays of HuCCT1, RBE, HCCC9810, and QBC939 cells treated with 100 ng/mL Thbs1 or none (NC). **(D)** CCK-8 assay measuring cell viability of HuCCT1, RBE, and QBC939 cells treated with 10 ng/mL Thbs1. Data are presented as mean ± SD. Significance was determined by a two-tailed unpaired t-test (*$p < 0.05$). **(E)** Representative Transwell migration chamber images of LV3-shCd47 knockdown cells cultured with BMDMs or complete medium in the lower chamber for 24 h. **(F)** Images of mice bearing subcutaneous allografts derived from BR cells infected with LV3-shNC or LV3-shCd47. **(G)** Quantification of the weight of subcutaneous allografts from mice injected with LV3-shNC or LV3-shCd47 infected BR cells. Data are presented as mean ± SD. Significance was determined by a two-tailed unpaired t-test (**$p < 0.01$, n = 6 mice). **(H)** Images of collected ascites from mice intraperitoneally injected with LV3-shNC (n = 3 mice) or LV3-shCd47 infected BR cells (n = 5 mice).

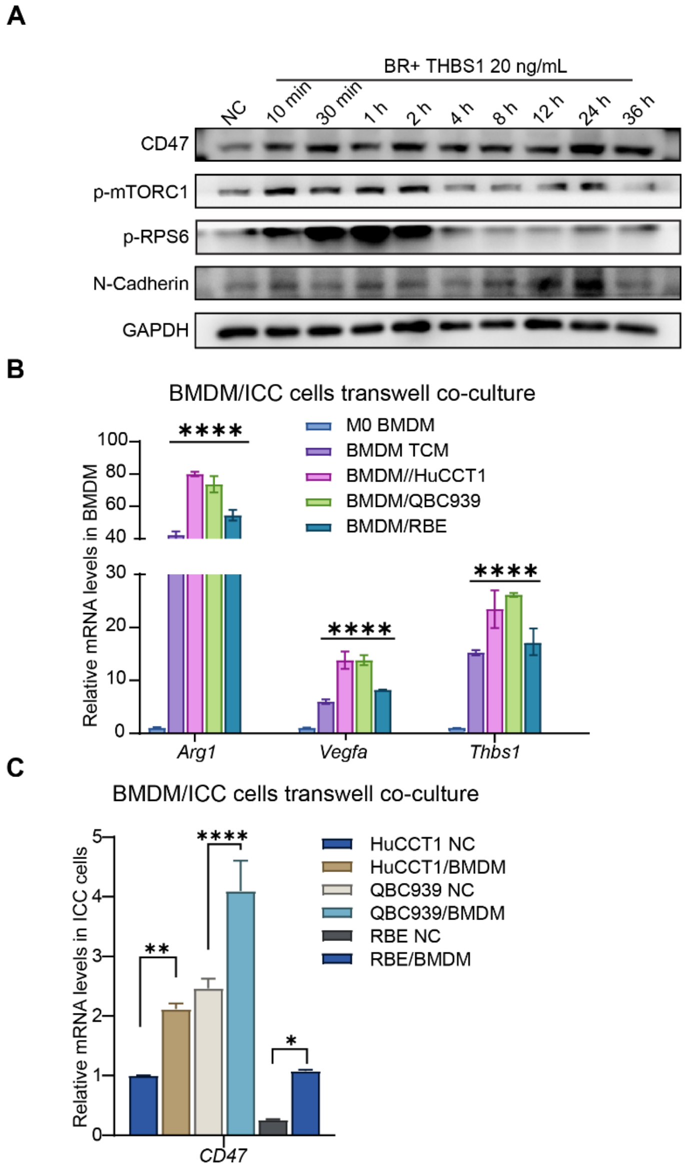

**Figure S10. Activated signaling or proteins in BR or HuCCT1 cells upon THBS1 treatment and BMDMs treated with TCM or co-cultured with ICC results.** (A) Western blot analysis of CD47, p-mTORC1 (Ser2448), p-RPS6, and N-Cadherinin BR cells stimulated with THBS1 over multiple time points. GAPDH served as the loading control. (B) qRT-PCR analysis of Arg1, Vegfa, and Thbs1 transcript levels in BMDMs treated with TCM or co-cultured with ICC cell lines in Transwell systems for 48 h. Data are presented as mean ± SD. Statistical significance was analyzed by one-way ANOVA ($p < 0.05$) followed by Dunnett's test (****$P < 0.0001$). (C) qRT-PCR analysis of CD47 transcript levels in ICC cells co-cultured with BMDMs using a Transwell system for 48 h. Data are presented as mean ± SD. Statistical significance was analyzed by one-way ANOVA ($p < 0.05$) followed by Dunnett's test (*$p < 0.05$, **$P < 0.01$, ****$P < 0.0001$, $n = 3$ independent experiments).

### Supplementary Tables

**Table S1 Organisms**

| Name | Supplier | Strain | Sex | Age | Overall n number |
|---|---|---|---|---|---|
| C57BL/6 mouse | Beijing Vital River LaboratoryAnimal Technology Co., Ltd | C57BL/6J | ♀/♂ | 6-8 w | 20 |
| FVB/N mouse | Beijing Vital River LaboratoryAnimal Technology Co., Ltd | FVB/NJ | ♀/♂ | 6-8 w | 30 |
| OSMR$^{fl/fl}$;Alb-Cre mouse | Shulaibao (Wuhan) Biotechnology Co., Ltd. | C57BL/6J | ♀/♂ | 6-8 w | 24 |
| OSMR$^{fl/fl}$ mouse | Shulaibao (Wuhan) Biotechnology Co., Ltd. | C57BL/6J | ♀/♂ | 6-8 w | 24 |
| Lifr$^{fl/fl}$;Alb-Cre mouse | Shulaibao (Wuhan) Biotechnology Co., Ltd. | C57BL/6J | ♀/♂ | 6-8 w | 6 |
| Lifr$^{fl/fl}$ mouse | Shulaibao (Wuhan) Biotechnology Co., Ltd. | C57BL/6J | ♀/♂ | 6-8 w | 6 |
| Thbs1$^{fl/fl}$;Lyz-Cre mouse | Shulaibao (Wuhan) Biotechnology Co., Ltd. | C57BL/6J | ♀/♂ | 6-8 w | 24 |
| Thbs1$^{fl/fl}$ mouse | Shulaibao (Wuhan) Biotechnology Co., Ltd. | C57BL/6J | ♀/♂ | 6-8 w | 24 |

**Table S2. Deposited data.**

| Name of repository | Identifier | Link | prognosis |
|---|---|---|---|
| scRNA-seq data | GSE308574 (This paper) | https://www.ncbi.nlm.nih.gov/geo/query/acc.cgi?acc= GSE308574 | No |
| Human ICC | GSE138709(2) | https://www.ncbi.nlm.nih.gov/geo/query/acc.cgi?acc=GSE138709 | No |

| | | | |
|---|---|---|---|
| scRNA-seq data | | | |
| Human ICC scRNA-seq data | GSE189903(Intrahepatic cholangiocarcinoma samples)(3) | https://www.ncbi.nlm.nih.gov/geo/query/acc.cgi?acc=GSE189903 | No |
| Human ICC scRNA-seq data | GSE210066(Intrahepatic cholangiocarcinoma)(4) | https://www.ncbi.nlm.nih.gov/geo/query/acc.cgi?acc=GSE210066 | No |
| Human ICC scRNA-seq data | GSE201425(CCA, primary focus)(5) | https://www.ncbi.nlm.nih.gov/geo/query/acc.cgi?acc=GSE201425 | No |
| ICC bulk RNA | E-MTAB-6389 | https://www.ebi.ac.uk/biostudies/arrayexpress/studies/E-MTAB-6389 | yes |
| TCGA(CHOL) | TCGA-CHOL[a] | (https://portal.gdc.cancer.gov) | Yes |
| ICC bulk RNA | GSE107943(6) | https://www.ncbi.nlm.nih.gov/geo/query/acc.cgi?acc=GSE107943 | Yes |
| ICC bulk RNA | GSE76297 | https://www.ncbi.nlm.nih.gov/geo/query/acc.cgi?acc=GSE76297 | No |
| ICC bulk RNA | GSE89749(7) | https://www.ncbi.nlm.nih.gov/geo/query/acc.cgi?acc=GSE89749 | Yes |
| ICC bulk RNA | GSE32225(8) | https://www.ncbi.nlm.nih.gov/geo/query/acc.cgi?acc=GSE32225 | No |
| ICC bulk RNA | GSE26566(9) | https://www.ncbi.nlm.nih.gov/geo/query/acc.cgi?acc= GSE26566 | No |
| ICC bulk RNA | GSE32879(10) | https://www.ncbi.nlm.nih.gov/geo/query/acc.cgi?acc=GSE32879 | No |
| ICC bulk RNA | GSE244807(10) | https://www.ncbi.nlm.nih.gov/geo/query/acc.cgi?acc= GSE244807 | Yes |
| spRNAseq | HRA000437(11) | https://ngdc.cncb.ac.cn/gsa-human/browse/HRA000437 | No |

**Table S3. Cell types and marker genes used for cell type definition.**

| Cell Type | Marker Genes |
|---|---|
| B | *CD19,CD79A,CD79B,IGHG1,IGHA* |

| | |
|---|---|
| Cholangiocyte | *KRT19,KRT7,EPCAM,SOX9,KRT18* |
| DC | *SIGLECH,XCR1,CD1C,FLT3,FCER1A,LAMP3,IDO1* |
| Endothelial | *ENG,KDR,PECAM1,LYVE1* |
| Fibroblast | *COL1A1,CoL1A2,DCN,ACTA2* |
| Hepatocyte | *ALB,HNF1B,CY2E1,CY2F2* |
| Macrophage | *ADGRE1,CCR2,CD68,CLEC4F* |
| Neutrophile | *NGP,CAMP,CSF3R* |
| NK | *NKG7,XCL1* |
| T | *CD3D,CD3E,CD28,TRAC* |

**Table S4 Mφ_VEGFA signatures**

| Cell Type | Marker Genes |
|---|---|
| Mφ_VEGFA | *GPR183,OLR1,EREG,CAPG,TREM2,BTG1,LYZ,PLAUR,ZNF331,CD9* |

**Table S5 OSMR protein interaction genes**

| Interaction genes |
|---|
| *Osmr,Ccnd1,Il1b,Vegfa,Egr1,Vim,Il6st,Irf1,Nfkb1,Hif1a,Tnf,Cdkn1a,Myc,Zfp36,Ubc,Gadd45b,Stat3,Fos,Epha2,Junb,Nav2,Jun,Dusp1,Atf3,Ier3,Icam1,Mcl1,Hes1,Klf6,Myh9,Pde4d,Odc1,Ier2,Tgif1,Zfp36l1,Lpp,Cyp1b1,Fosb* |

**Table S6 Antibodies**

| Product Name | Target | Supplier | Catalog | Dilution |
|---|---|---|---|---|

| | | | Number | |
|---|---|---|---|---|
| CD206/MRC1 (E6T5J) XP® Rabbit mAb | CD206 | CST | 24595S | WB 1:2000; IHC 1:500; mIF 1:500 |
| Vimentin Polyclonal antibody | Vimentin | proteintech | 10366-1-AP | WB 1:20000 |
| Phospho-mTOR (Ser2448) Antibody | p-mTORC1 | CST | 2971S | WB 1:2000 |
| Alpha Tubulin Monoclonal antibody | α-tubulin | proteintech | 66031-1-Ig | WB 1:20000 |
| APC anti-mouse CD45 Antibody(30-F11) | CD45 | BioLegend | 103112 | FC 1:100 |
| PerCP/Cyanine5.5 anti-mouse F4/80 Recombinant Antibody(QA17A29) | F4/80 | BioLegend | 157318 | FC 1:100 |
| FITC anti-mouse/human CD11b Antibody(M1/70) | CD11b | BioLegend | 101206 | FC 1:100 |
| APC anti-mouse CD206 (MMR) Antibody(C068C2) | CD206 | BioLegend | 141708 | FC 1:100 |
| OSMR Recombinant monoclonal antibody | OSMR | proteintech | 84555-2-RR | mIF 1:500 |
| Anti-OSMR antibody [EPR28222-64] | OSMR | Abcam | ab315388 | WB 1:2000 |
| Cyclin D1 Monoclonal antibody | CCND1 | proteintech | 60186-1-Ig | WB 1:2000 |
| BCL2 Monoclonal antibody | BCL2 | proteintech | 60178-1-Ig | WB 1:2000 |
| Phospho-Stat3 (Tyr705) (D3A7) XP® Rabbit mAb | p-Stat3 | CST | 9145T | WB 1:2000 |
| CD47 Recombinant monoclonal antibody | CD47 | proteintech | 85958-1-RR | WB 1:2000 |
| N-Cadherin (D4R1H) XP® Rabbit mAb | N-Cadherin | CST | 13116S | WB 1:2000 |
| Anti-phospho Drebrin(Ser142) | p-DBN1 | merck | MABN833 | WB 1:1000 |
| DBN1 Antibody | DBN1 | CST | 96540S | WB 1:2000 |
| Phospho-S6 Ribosomal protein | p-RPS6 | proteintech | 29223-1-AP | WB 1:2000 |

| | | | | |
|---|---|---|---|---|
| (Ser235/236) Polyclonal antibody | | | | |
| Cytokeratin 19 Polyclonal antibody | CK19 | proteintech | 10712-1-AP | IHC 1:200; mIF 1:500 |
| Ki-67 Polyclonal antibody | Ki67 | proteintech | 27309-1-AP | IHC 1:200; mIF 1:1000 |
| F4/80 Polyclonal antibody | F4/80 | proteintech | 28463-1-AP | IHC 1:800 |
| Oncostatin M Polyclonal antibody | OSM | proteintech | 27792-1-AP | WB 1:2000; IHC 1:500; mIF 1:300 |
| Thrombospondin 1 Polyclonal antibody | THBS1 | proteintech | 18304-1-AP | WB 1:2000; mIF 1:200 |
| Osteopontin Polyclonal antibody | SPP1 | proteintech | 22952-1-AP | WB 1:2000; IHC 1:200; mIF 1:500 |
| Anti-CD68 antibody (IHC) | CD68 | AiFang biological | AF20022 | mIF 1:600 |
| Goat Anti-Mouse IgG H&L（HRP） | Anti-Mouse IgG | Zenbio | 511103 | WB 1:2000 |
| Rabbit Anti-Goat IgG H&L (HRP) | Anti-Rabbit IgG | Zenbio | 550094 | WB 1:2000 |

**Table S7 qRT-PCR primers**

| **Gene** | **species** | **Forward primer (5'-3')** | **Reverse primer (5'-3')** |
|---|---|---|---|
| *Actb* | Mouse | GGCTGTATTCCCCTCCATCG | CCAGTTGGTAACAATGCCATGT |
| *Vegfa* | Mouse | CCACGACAGAAGGAGAGCAGAAGTCC | CGTTACAGCAGCCTGCACAGCG |
| *SPP1* | Mouse | AGCAAGAAACTCTTCCAAGCAA | GTGAGATTCGTCAGATTCATCCG |
| *Thbs1* | Mouse | GGGGAGATAACGGTGTGTTTG | CGGGGATCAGGTTGGCATT |
| *Tgfbr2* | Mouse | TTGGATTGCCAGTGCTAACCC | AACAAGCCACAGTAACATGACA |
| *Osmr* | Mouse | CATCCCGAAGCGAAGTCTTGG | GGCTGGGACAGTCCATTCTAAA |
| *OSMR* | Human | AATGTCAGTGAAGGCATGAAAGG | GAAGGTTGTTTAGACCACCCC |
| *Osm* | Mouse | CAGAATCAGGCGAACCTCACG | AGCTCTCAGGTCAGGTGTGTT |
| *Lifr* | Mouse | AGCTCTGACCCTCCTGCAT | TGGGTGACAAGAATGGAACCT |
| *IL10* | Mouse | GCTCTTACTGACTGGCATGAG | CGCAGCTCTAGGAGCATGTG |
| *Cd47* | Mouse | TGCGGTTCAGCTCAACTACTG | GCTTTGCGCCTCCACATTAC |
| *CD47* | Human | TCCGGTGGTATGGATGAGAAA | ACCAAGGCCAGTAGCATTCTT |
| *Arg1* | Mouse | CCACAGTCTGGCAGTTGGAAG | GGTTGTCAGGGGAGTGTTGATG |
| *ACTB* | Human | CATGTACGTTGCTATCCAGGC | CTCCTTAATGTCACGCACGAT |

**Table S8 siRNA sequence**

| siRNA Name | Target Species | Strand Type | sequence（5'→3'） |
|---|---|---|---|
| siCD47-1 | Human | Sense | AAGUCACAAUUAAACCAAGGCTT |
| | | Antisense | GCCUUGGUUUAAUUGUGACUUTT |
| siCD47-2 | Human | Sense | AACUAGUCCAAGUAAUUGUGCTT |
| | | Antisense | GCACAAUUACUUGGACUAGUUTT |
| siOSMR-402 | Human | Sense | AUGUUGUCUGAAAGACUGCTT |
| | | Antisense | GCAGUCUUUCAGACAACAUTT |
| siOSMR-863 | Human | Sense | AACAUUGGUGCCUUCUUCCTT |
| | | Antisense | GGAAGAAGGCACCAAUGUUTT |
| siOSMR-1200 | Human | Sense | AAGUGUAGCUUUGGGAAGGTT |
| | | Antisense | CCUUCCCAAAGCUACACUUTT |
| siScramble | Universal | Sense | ACGUGACACGUUCGGAGAATT |
| | | Antisense | UUCUCCGAACGUGUCACGUTT |

**Table S9 shRNA and sgRNA sequence**

| Name | Target Gene | Target species | Vehicle | Sequence（5'→3'） |
|---|---|---|---|---|
| LV3-shCD47 | CD47 | Mouse | LV3 (H1/GFP&Puro) | CAGTCTCAGACTTAATCAA |
| LV3-shNC | NC | Mouse | LV3 (H1/GFP&Puro) | ACTACCGTTGTTATAGGTG |
| sgOSMR-1 | OSMR | Human | pLenti-CRISPR-v2 | GAACATACCAGCTTCAAGTG |
| sgOSMR-2 | OSMR | Human | pLenti-CRISPR-v2 | TACTCGCGCCATGTACTCTG |
| sgOSMR-3 | OSMR | Human | pLenti-CRISPR-v2 | AGGGACAAATATCTATTGTG |
| sgOSMR-4 | OSMR | Huma | pLenti-CRISPR- | GTAAGTGTGCAAATTCTCTG |

| | | | | |
|---|---|---|---|---|
| | | n | v2 | TGG |

**Table S10 Reagents, inhibitors and recombinant proteins**

| **Reagents, inhibitors and recombinant proteins** | **supplier** | **Catalog Number** |
|---|---|---|
| Clophosome® and Control Liposomes | FormuMax | F70101C-NC |
| Plain Control Liposomes for Clophosome® (Neutral) | FormuMax | F70101C-N |
| BCA Protein Colorimetric Assay Kit | Elabscience | E-BC-K318-M |
| Protease Inhibitor Cocktail (EDTA-Free, 100× in DMSO) | MCE | HY-K0010 |
| RIPA Lysis Buffer (Strong) | MCE | HY-K1001 |
| PMSF | MCE | HY-B0496 |
| Tween-20 | BioFroxx | 1247ML100 |
| Triton X-100 | BioFroxx | 1139ML100 |
| SDS | BioFroxx | 3250GR500 |
| 2×TSINGKE® Master SYBR Green I qPCR Mix-UDG (Without ROX) | Tsingke | TSE204 |
| Endo-Free Plasmid DNA Maxi Kit | Omega | D6926-04 |
| OSM recombinant protein (Human) | R&D system | 295-OM-010 |
| THBS1 recombinant protein (Human) | MCE | HY-P70725 |
| Goldenstar® RT6 cDNA Synthesis Kit | Tsingke | TSK302S |
| BeyoClick™ EdU-555 Cell Proliferation Detection Kit<br>Rapamycin<br>BTP2 | Beyotime<br>MCE<br>MCE | C0075S<br>HY-10219<br>YM-58483 |
| SMI-OSM-10B | MCE | HY-148692 |

**Table S11 Software**

| Software name | Manufacturer | Version |
|---|---|---|
| Seurat | Hao et al[1] | V 4.4.0 |
| SciBet | Li et al[2] | V 1.0 |
| harmony | Korsunsky el al[3] | V 1.0 |
| BSTMatrix | http://www.biomarker.com.cn/zhizao/tools | V2.3 |
| Cottrazm | Xun et al[4] | V0.1.1 |
| STutility | https://ludvigla.github.io/STUtility_web_site/index.html | V1.1.1 |
| clusterProfiler | Yu et al[5] | V4.10 |
| spacexr | Cable et al[6] | V2.2.1 |
| xCell | Aran et al[7] | V 1.1.0 |
| infercnv | Puram et al[8] | V1.0.4 |
| Scissor | Sun et al[13] | V2.0.0 |
| CellphoneDB | Garcia et al[14] | V3 |
| CellChat | Jin et al[15] | V1.5 |
| NicheNet | Browaeys et al[16] | V2.1.0 |
| Monocle | Trapnell et al[17] | V2.14.0 |
| ArchR | Granja et al[18] | V1.0.3 |
